# Functional Specialization of S-Adenosylmethionine Synthases Links Phosphatidylcholine to Mitochondrial Function and Stress Survival

**DOI:** 10.1101/2025.02.20.639242

**Authors:** Athena L. Munden, Dominique S. Lui, Daniel P. Higgins, Matthew J. Fanelli, Thien-Kim Ngyuen, Katherine M. Edwards, Maria Ericsson, Adwait A. Godbole, John A. Haley, Caroline Lewis, Jessica B. Spinelli, Benjamin Harrison, Daniel Raftery, Danijel Djukovic, Daniel E.L. Promislow, Dana L. Miller, Amy K. Walker

## Abstract

S-adenosylmethionine (SAM), produced by SAM synthases, is critical for various cellular regulatory pathways and the synthesis of diverse metabolites. Studies have often equated the effects of knocking down one synthase with broader SAM-dependent outcomes such as histone methylation or phosphatidylcholine (PC) production. Humans and many other organisms express multiple SAM synthases. Evidence in *Caenorhabditis elegans*, which possesses four SAM synthase genes, suggest that the enzymatic source of SAM impacts its function. For instance, loss of *sams-1* leads to enhanced heat shock survival and increased lifespan, whereas reducing *sams-4* adversely affects heat stress survival. Here, we show that SAMS-1 contributes to a variety of intermediary metabolic pathways, whereas SAMS-4 is more important to generate SAM for methylation reactions. We demonstrate that loss of *sams-1* exerts age-dependent effects on nuclear-encoded mitochondrial gene expression, mitochondrial metabolites, and may induce mitophagy. We propose a mechanistic model where reduced SAM from SAMS-1 acts through PC to impact mitochondria, thereby enhancing survival during heat stress.

## Introduction

Remodeling gene expression to adapt to distinct physiological conditions is a common strategy for countering exogenous stressors (*1*). The changes in gene expression depend on the nature of the stressor; defensive peptides are produced in response to pathogen stress, detoxification enzymes are upregulated in oxidative stress, and chaperones are expressed upon protein folding stress. The metabolic state of the cells also influences the transcriptional response to stress (*2*). For example, limitation of the 1-carbon cycle metabolite SAM (S-adenosylmethionine) has distinct effects on stress responses, depending on the type of stress and the metabolic source of SAM (*3–5*).

SAM could influence gene expression through its role as the predominant cellular methyl donor, linking histone methylation and chromatin regulatory patterns. However, SAM also contributes to other metabolic pathways that generate phosphatidylcholine, glutathione, and or polyamines. Changes in these pathways could also have indirect effects on gene expression (*6*). This pleiotropy underscores the need to identify mechanistic links between SAM, its downstream metabolites or methylation targets, and phenotypes resulting from SAM depletion across phyla, such as altered stress responses, lipid accumulation, changes in differentiation or development, and extended lifespan (*3*, *7–12*).

SAM is produced from methionine and ATP by SAM synthases, a highly conserved enzyme present as multiple paralogs in many eukaryotes. Yeast and mammals both have two synthases (SAM1/SAM2 and MAT1A/MAT2A, respectively). In yeast, the synthases have distinct effects on gene expression and genome stability (*13*). In mammals, SAM synthases are expressed in different tissues and have distinct roles. MAT1A is specific to adult liver and KO mice have fatty liver while MAT2A is expressed widely (*14*). MAT2A also has a regulatory partner, MAT2B, which can form multiple heteromeric complexes with distinct enzymatic characteristics (*14*). However, understanding the mechanistic properties of these synthases or isoforms have been difficult to study in mammals, as MAT1A expression decreases in cultured cells, MAT2A is essential for viability, and cell culture media is replete with 1-carbon cycle (1CC) intermediates (*15*). To circumvent this issue, we have focused on SAM synthases in *C. elegans*, where the family has been extended to 4 paralogs, *sams-1 (X)*, the bicistronically expressed *sams-3 (IV), sams-4 (IV)*, and *sams-5 (IV)*. While germline tissue express a subset of the synthases, most somatic cells express at least two (*5*) and knockdown of individual synthases have similar effects on total SAM levels (*3*, *5*, *7*). However, phenotypes after SAM synthase knockdown are dramatically different.

Animals lacking *sams-1* have increased lipid stores (*7*), long lifespan (*8*), are sensitive to bacterial stress (*3*), and live longer after heat shock (*5*). In contrast, reduction in *sams-4* increases sensitivity to heat shock but has negligible effects on lifespan (*5*), suggesting that each SAMS is connected to distinct downstream metabolic products or methylation patterns. Global H3K4me3 patterns change after RNAi of either *sams-1* or *sams-4*, but RNAi of the H3K4 methyltransferase *set-16*/MLL phenocopied the heat sensitivity of *sams-4* mutant animals (*4*, *5*). Production of SAM and H3K4me3 levels increased after heat shock in *sams-1(lof)* animals in a *sams-4*-dependent fashion (*5*), linking production of SAM from *sams-4* (SAM_4_) to chromatin regulatory patterns necessary for heat stress survival. The link between the loss of SAM from *sams-1* (SAM_1_) and enhanced survival was less clear, as *sams-1* animals exhibit changes in both histone methylation patterns and lipid production.

In this study, we show that SAM_1_ is required for a variety of mitochondrial metabolites, expression of mitochondrial encoded genes during aging, and mitophagy. We further demonstrate that these effects on mitochondria the requirement for SAM_1_ activity for production of PC, a crucial mitochondrial membrane lipid. Collectively, these results further support the idea that individual SAMS enzymes provision specific methylation pathways with independent physiological effects.

## Results

### SAMS-1, but not SAMS-4, affects a variety of cellular metabolites

The *sams-1* and *sams-4* genes both encode SAM synthases (Fig 1A, see also Fig 2A for a Methionine/SAM cycle diagram). However, depletion of *sams-1* and *sams-4* results in different phenotypes, suggesting that SAM produced by each enzyme may function in different cellular processes or integrate with distinct metabolic pathways. To explore this possibility, we performed targeted metabolomics in animals lacking SAM production by either SAMS-1 or SAMS-4. We focused initially on the effect of heat shock because the loss of *sams-1* or *sams-4* have opposite effects. We identified 62 metabolites that significantly differed across conditions (Table S1). Fig. S1 shows the relationships between selected metabolites using pathway diagrams based on WormPaths (*16*).

**Figure 1:**
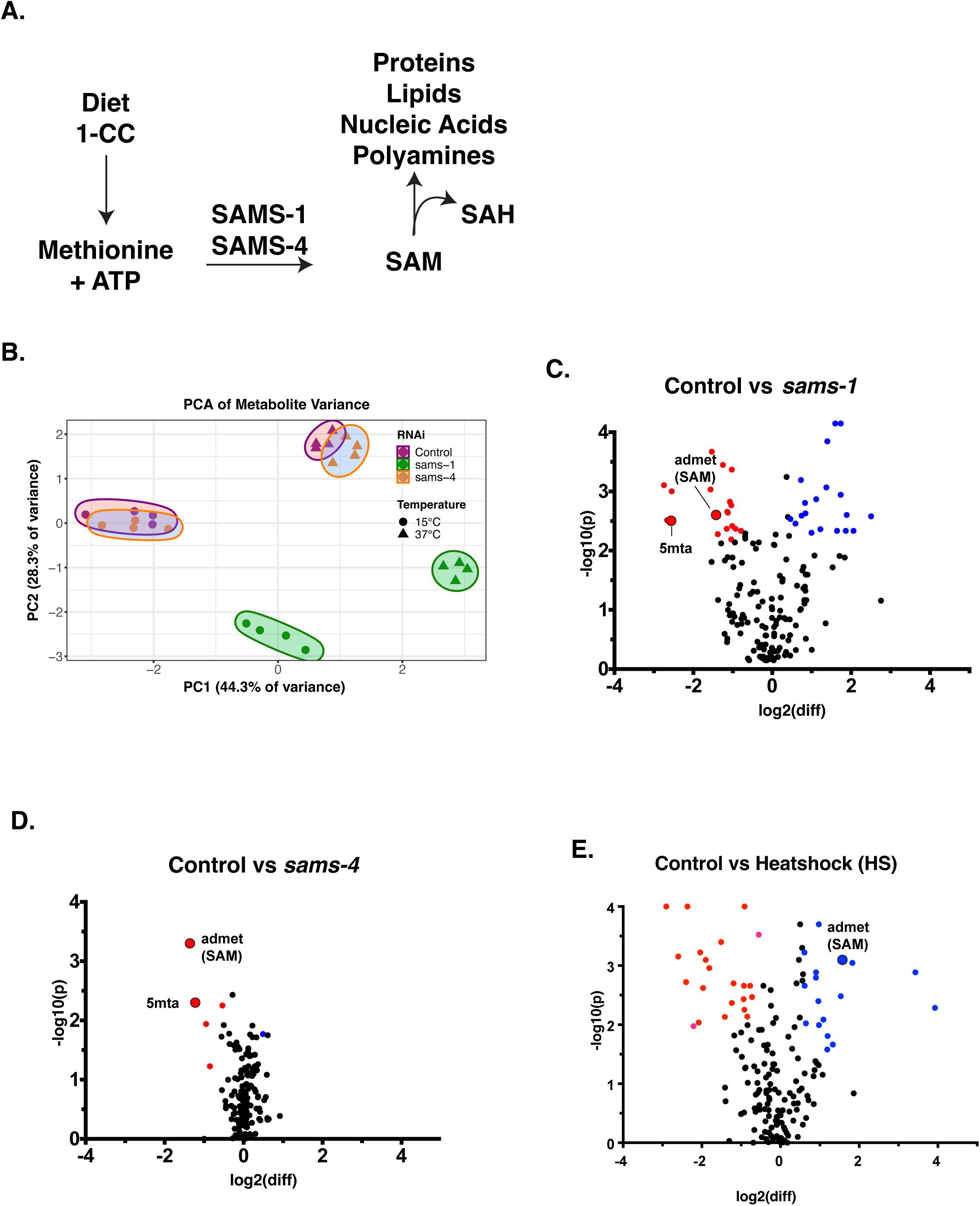
Targeted metabolomics shows broad metabolic changes in *sams-1* animals in basal and heat shock conditions. **A.** Schematic of metabolic pathways using SAM. **B**. PCA chart demonstrating distinct components in *sams-1* animals at control and heat shock conditions. Volcano plots comparing log10 fold changes in metabolite levels between Control and *sams-1(RNAi)* (**C.**) or Control and *sams-4* (**D.**) samples and Control vs Heatshock (**E.**). 1-carbon cycle metabolites admet (SAM) and 5mta (methyladenosine) are highlighted.

**Figure 2:**
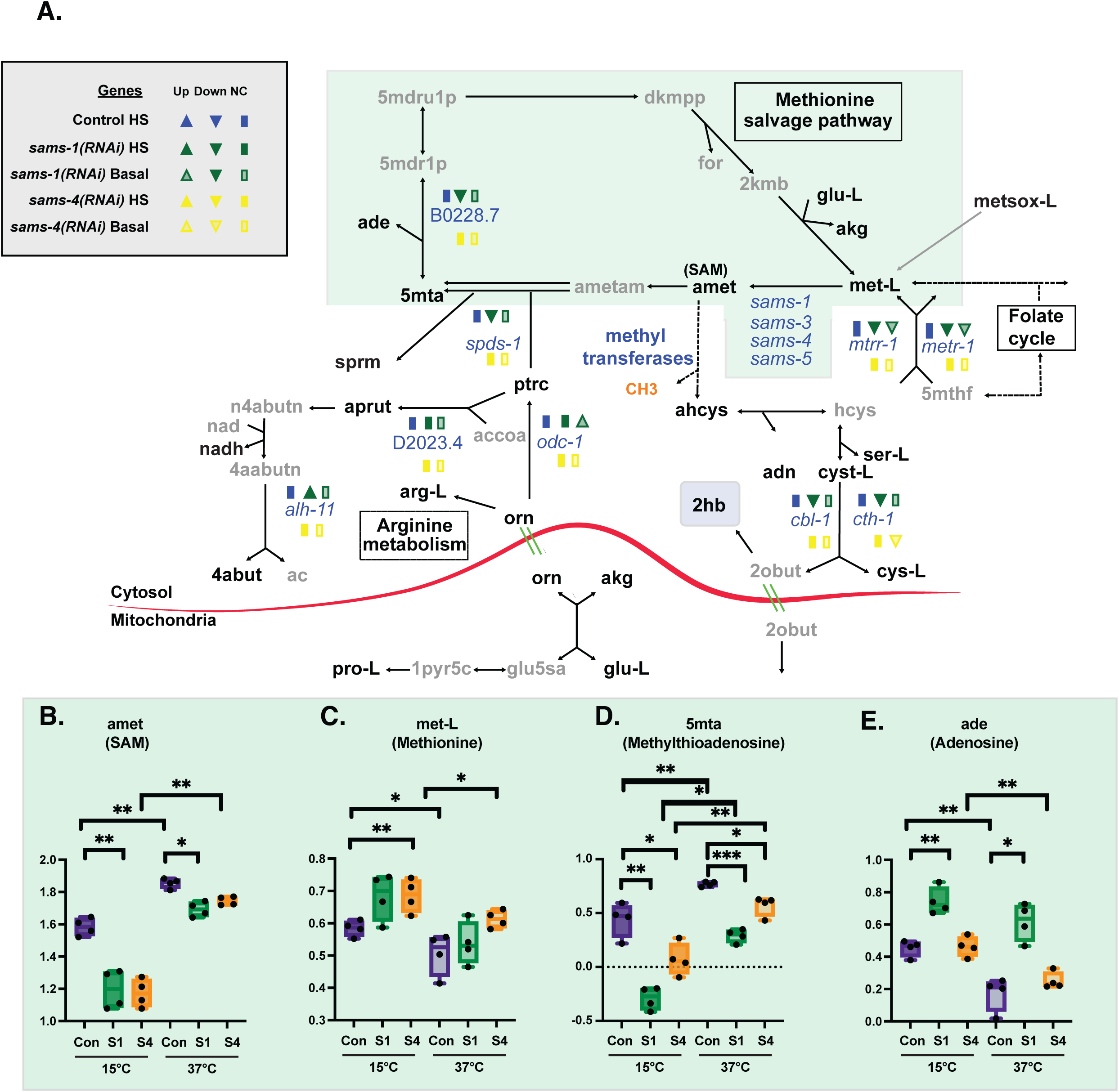
Distinct effects on methionine salvage and polyamine pathways in sams-1 and sams-4(RNAi) animals. **A**. Schematic derived from WormPaths (*16*) showing methionine salvage and arginine derived metabolic pathways. See **Figure S1** for schematic linking these pathways to mitochondrial fatty acid oxidation. Information regarding gene level and H3K4me3 status from Control, *sams-1* or *sams-4* animals in basal or heat shocked conditions is marked in legend. Data is from Godbole, et al. eLife 2023. Levels of selected metabolites from methionine salvage (green box) (**B-E**) are shown by box and whisker plots. Whiskers encompass data range and significance is determined by two-way repeated measures ANOVA. ns: q-value ≥ 0.05 *: q-value <0.05, **: q-value <0.01, ***: q-value <.001, ****: q-value > 0.0001.

Metabolites will be referred to with the same abbreviation used in WormPaths throughout this manuscript.

In unstressed animals, principal component analysis (PCA) shows widespread changes in metabolites in the *sams-1(lof)* animals, whereas the *sams-4(lof)* metabolite profile was quite similar to controls (Fig 1B, 15°C). The changes we observe in metabolite profiles resemble the differential effect of *sams-1* and *sams-4* on gene expression.

Widespread changes occurred after *sams-1(RNAi)*, whereas *sams-4(RNAi)* animals were similar to controls (*5*).

We next analyzed the metabolite profiles of animals exposed to heat stress, where *sams-4(lof)* die rapidly but *sams-1(lof)* survive better than wild-type (*5*). We identified 46 metabolites that significantly changed in animals after heat stress. PCA shows dramatic changes upon heat stress, but the pattern was the same as in basal conditions: the metabolite profile of *sams-4(RNAi)* animals after heat stress was similar to controls, whereas we observed broad and distinct changes in *sams-1(RNAi)* animals (Fig 1B, 37°C). The limited effects of *sams-4(RNAi)* on metabolite abundances supports the assertion that its role in histone methylation underlies the decreased survival of *sams-4(lof)* animals in heat stress. Further supporting this idea, the changes in gene expression we observe in *sams-4(RNAi)* animals exposed to heat stress (*5*) mirror changes when the histone methyltransferase *set-16* is depleted (*4*), suggesting that these changes are related to histone methylation. In contrast, our data indicate that increased survival of *sams-1(lof)* animals in heat stress is associated with changes beyond gene expression.

To define the specific effects of *sams-1* and *sams-4*, we identified the metabolites that changed when each SAMS enzyme was depleted. Consistent with our PCA, we observed many changes in metabolite abundance in *sams-1(RNAi)* animals in basal conditions (Fig 1C). In contrast, only a few metabolites decreased in *sams-4(RNAi)* animals (Fig 1D). Importantly, we found that steady-state SAM (amet) abundance decreased by ∼50% after RNAi of either *sams-1* or *sams-4* (Fig 2B), consistent with previous results (*3*, *5*, *7*). These data demonstrate that the differences we observe between *sams-1* and *sams-4* animals do not result from quantitative changes in SAM production. Moreover, the fact that depletion of either SAMS enzyme reduces the abundance of SAM indicates that these enzymes do not compensate for each other in basal conditions.

### SAM from SAMS-1 and SAMS-4 contributes to different metabolic pathways in heat stress

When control animals are exposed to heat stress there is a widespread change in metabolite profile (Fig 1E). Consistent with previous reports (*5*), SAM (amet) was one of the metabolites that was more abundant after heat stress in wildtype animals (Fig 1E). We noticed a concomitant decrease in methionine levels in heat-stressed animals (Fig 2C), which could reflect higher SAMS enzyme activity in these conditions. Notably, SAM increased dramatically in both *sams-1* and *sams-4(RNAi)* animals exposed to high temperature (Fig 2B), indicating that neither enzyme is uniquely required for SAM production in heat stress. These two enzymes may compensate for each other in heat stress (but not in basal conditions); alternatively, increased activity of other SAMS enzymes might generate SAM in heat stress.

Under basal conditions, depletion of *sams-1* and *sams-4* both impacted 1CC-related pathways - but with some notable differences. For example, S-adenosyl homocysteine (ahcys) was less abundant specifically in *sams-4(RNAi)* animals, which could suggest a decrease in SAM-dependent methylation reactions (Fig S2A). In contrast, levels of 5-methylthioadenosine (5mta) and adenosine (ade) were perturbed only in *sams-1(RNAi)* animals (Fig 2D, 1E). 5-methylthioadenosine (5mta) is generated when SAM is used in polyamine synthesis (*17*), and adenosine (ade) is released in this methionine salvage pathway. In *sams-1(RNAi)* animals under both basal and heat-stress conditions, 5-methylthioadenosine (5mta) is low and adenosine (ade) is high (Fig 2D,E), indicative of increased methionine salvage. Our data also are consistent with increased polyamine synthesis in *sams-1(RNAi)* animals: we found increased levels of polyamines such as putrescine (ptrc) and N-acetylputrescine (apurt) specifically in *sams-1(RNAi)* animals (Fig S2B,C), and in heat stress the abundance of ornithine (orn), a substrate for polyamine synthesis, is particularly sensitive to loss of *sams-1(RNAi)* (Fig S2D). These data suggest that *sams-1* is important to coordinate at least two metabolic processes, methionine salvage and polyamine synthesis, whereas *sams-4* may be more important for methyltransferase activity.

Depletion of *sams-1 or sams-4* also had different effects on the transsulfuration pathway (Fig 2A), which produces cysteine (cys-L) from homocysteine (hcys). During heat stress, even wild-type animals have dramatically lower levels of 2-hydroxybutyrate (2hb), which is produced from the cleavage of cystathionine (cyst-L) to generate cysteine (cys-L), suggesting alterations in the transsulfuration pathway. This result suggests that transsulfuration is reduced even as levels of SAM (amet), a positive regulator of transsulfuration by allosteric regulation of cystathionine β-synthase (*18*), increase (Fig 2B). RNAi of either *sams* gene in basal conditions results in similar decreases to each other in abundance of 2-hydroxybutyrate (2hb) (Fig S2E), possibly indicating a reduction in transsulfuration activity reflecting a reduced flux through the methionine cycle. In contrast, heat stress increases the level of 2-hydroxybutyrate (2hb) in *sams-1(RNAi)* animals and does not change it in *sams-4(RNAi)* animals (Fig S2E).

This suggests that *sams-1* in particular is important to modulate flux through the transsulfuration pathway during heat stress.

### SAMS-1 promotes fatty acid oxidation and detoxification of propionyl-CoA

To better understand how loss of *sams-1* perturbs cellular metabolism, we mapped the metabolites that changed in *sams-1(lof)* animals onto WormPaths (*16*). In addition to changes in methionine metabolism noted above (Fig 2), we found that *sams-1(RNAi)* had specific effects on metabolites associated with fatty acid β-oxidation (Fig 3A). In basal conditions, carnitine (crn), which is used to transport very long chain fatty acids into the mitochondria for β-oxidation, was elevated in *sams-1(RNAi)*, but not *sams-4(RNAi),* animals (SFig2G). Heat stress resulted in a slight increase in carnitine in wild-type and *sams-4(RNAi)* animals, but not to the level of accumulation observed in *sams-1(RNAi)* animals. These data suggest that β-oxidation is perturbed in the *sams-1(lof)* animals. It may be that this change is not a direct result of reduced SAM (amet) production by SAMS-1, as the expression of several carnitine palmitoyl transferase (cpt) enzymes are lower in *sams-1(RNAi)* animals (Fig 3A, *5*).

**Figure 3:**
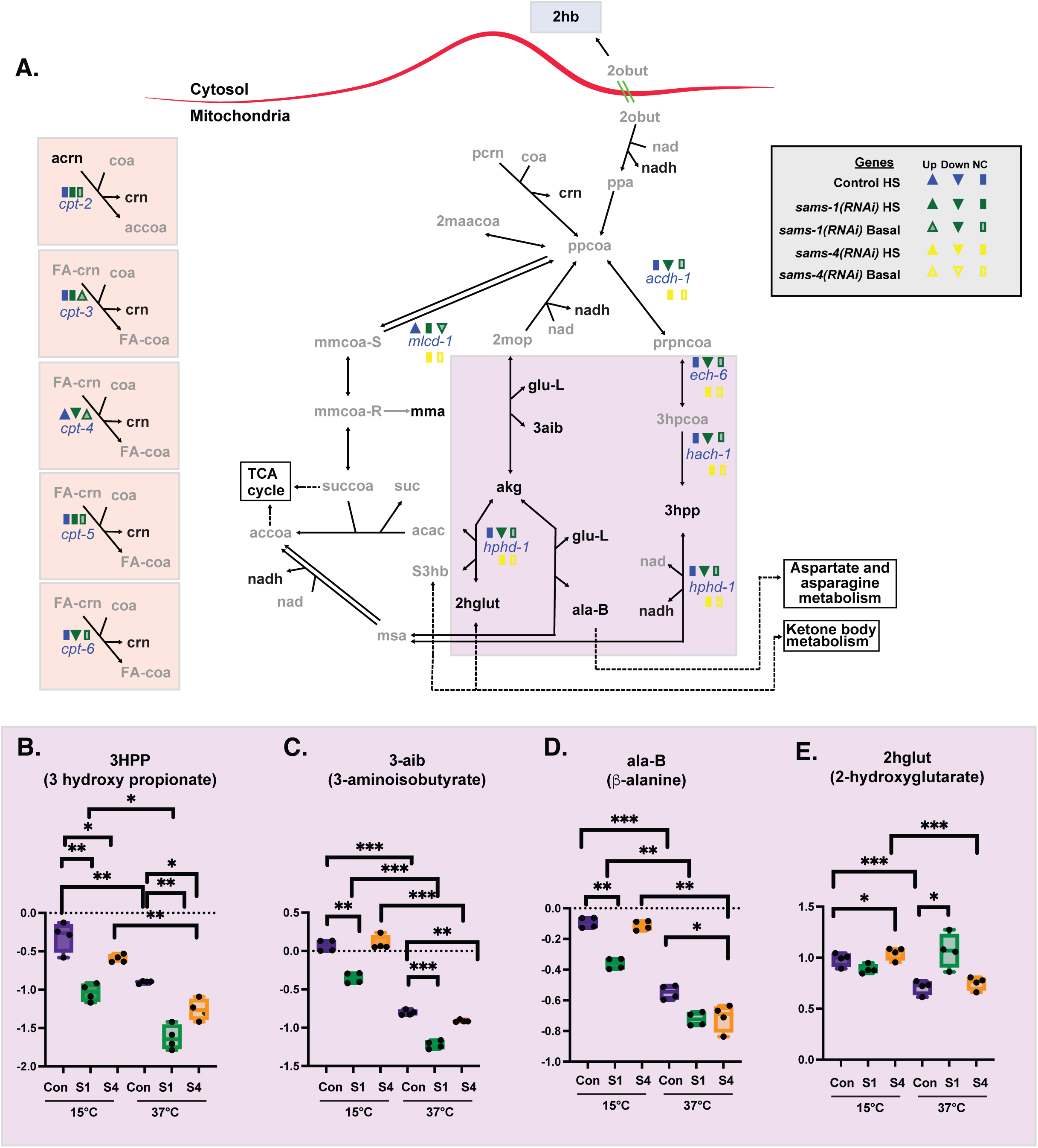
Heat shock accentuates changes in multiple mitochondrial metabolites after sams-1(RNAi). **A**. Schematic derived from WormPaths (*16*) showing methionine salvage and arginine derived metabolic pathways. See Figure S2 for schematic linking these pathways to methionine salvage and arginine/polyamine synthesis pathways. Information regarding gene expression level and H3K4me3 status from Control, *sams-1* or *sams-4* animals in basal or heat shocked conditions is marked in legend, data is from (*5*). Levels of selected metabolites from are shown by box and whisker plot and linked by color box to specific regions of the schematic (**B-E)**. Whiskers encompass data range and significance is determined by two-way repeated measures ANOVA. ns: q-value ≥ 0.05 *: q-value <0.05, **: q-value <0.01, ***: q-value <.001, ****: q-value > 0.0001.

The abundance of several metabolites in the degradation of propionyl-CoA (ppcoa), a mitochondrial β-oxidation-like process, was also particularly sensitive to loss of *sams-1* (Fig3A). In basal conditions, levels of 3-hydroxypropionate (3hpp) and 3-aminoisobutyrate (3aib) were decreased in *sams-1(RNAi)* animals (Fig 3B,C). When heat-stressed, 3-hydroxypropionate (3hpp) and 3-aminoisobutyrate (3aib) levels decline in all strains, exacerbating the depletion in the *sams-1(RNAi)* animals. We observed a similar *sams-1*-associated depletion of β-alanine (ala-B) that was enhanced by heat-stress (Fig 3D). The one exception was the metabolite 2-hydroxyglutarate (2hglut), which was not depleted in *sams-1(RNAi)* animals in basal conditions (Fig 3E). With heat stress, 2-hydroxyglutarate (2hglut) levels decline in wild-type, similar to other metabolites in this pathway. However, in *sams-1(RNAi)* animals exposed to heat stress, the levels of 2-hydroxyglutarate (2hglut) were the same as in basal conditions (Fig 3E). It may be that compensation by other metabolic pathway, such as isocitrate or ketone body metabolism, contribute to the steady state pools more in these conditions.

Together, these data show that *sams-1* is required for proper catabolism of propionyl-CoA. Consistent with this interpretation, the expression of several genes that encode for enzymes in this pathway is reduced in *sams-1(RNAi)* animals (Fig 3A, (*5*)), suggesting reduction in both gene and expression and metabolites in this mitochondrial process.

Together, our metabolite profiling suggests that SAM produced by SAMS-1 has broad contributions to intermediary metabolic pathways, whereas SAM produced by SAMS-4 may be used preferentially as a methyl donor.

### SAMS-1 is required to maintain age-associated expression of mitochondrial genes

We reasoned that the decrease in metabolites from multiple mitochondrial pathways after *sams-1* RNAi might be linked to decreased expression of mitochondrial genes. To understand how gene expression might change in *sams-1* animals as they age, we measured mRNA abundance in WT and *sams-1(lof)* animals at day 1 (D1) and day 7 (D7) of adulthood by RNAseq (Fig 4A, Table S2). Since heat shock and lifespan increase can share mechanistic links, we performed a detailed analysis of this data.

**Figure 4:**
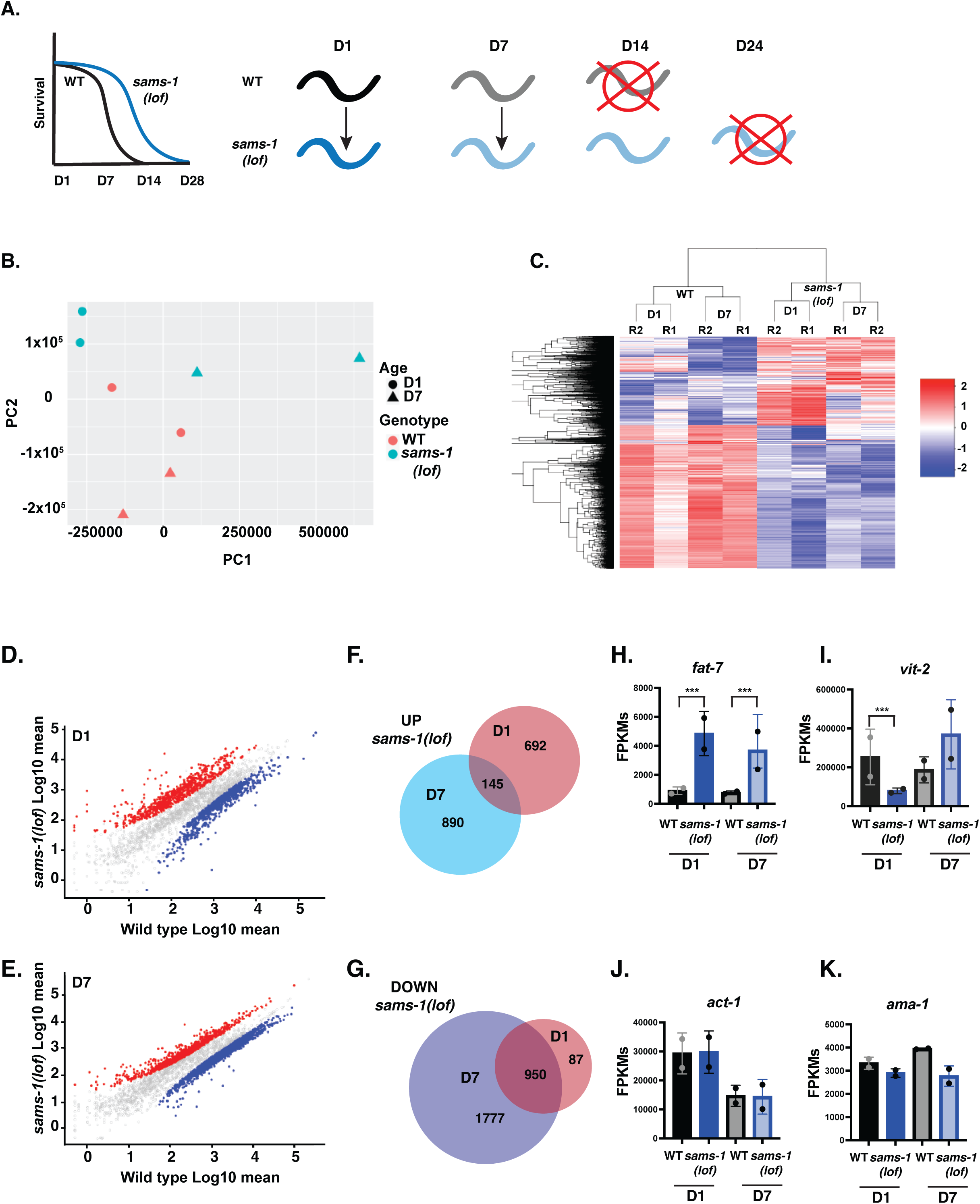
Aging *sams-1(lof)* animals express distinct sets of genes from wild type animals as they age. **A.** Schematic showing design of RNA seq experiment comparing younger and aged populations of long-lived *sams-1(lof)* animals. **B**. Principal component analysis shows separation of WT and *sams-1(lof)* animals as they age for all genes. **C**. Heatmap using k-means clustering demonstrating that gene expression patterns in wild type and *sams-1(lof)* animals cluster by genotype, rather than age. Scatter plots showing distribution of UP and DOWN regulated genes at D1 (**D**) and D7 (**E**). Venn Diagram comparing overlap between genes up (**F**) or down (**G**) regulated at D1 vs D7 in *sams-1(lof)* animals. Column graphs of FPKMs show two genes previously shown to be up or down regulated after *sams-1(RNAi) fat-7* (**H**) and *vit-2* (**I**) (*3*) as well as two housekeeping genes, *act-1* (b-actin) and *ama-1* (RNA Pol II beta subunit) (**J, K)**. Whiskers encompass standard deviation and the padjust value calculated by Deseq2 shows significance including a false discovery rate with * p <0.01, ** p <0.005, *** p 0.001.

PCA indicates that gene expression in *sams-1(lof)* animals is distinct from the wild-type control at both D1 and D7 (Fig 4B). This result is corroborated by clustering analysis (Fig 4C), which shows that changes in transcripts are more similar in *sams-1(lof)* across age (D1 vs D7) than with wild-type controls of the same age. Scatter plots showed more variation between *sams-1(lof)* and wild-type controls at D1 than at D7 (Fig 4, D, E), and relatively few of the genes that are upregulated in D1 *sams-1(lof)* animals are also upregulated at D7 (Fig 4F). In contrast, many genes downregulated at D1 in *sams-1(lof)* are further decreased in older animals at D7 (Fig4G). This suggests that in addition to a core subset of genes that are downregulated with aging, *sams-1(lof)* animals acquire a distinctive gene expression signature. Importantly, our data set was consistent with previous studies (*3*) where loss of *sams-1* induced the upregulation of *fat-7* and decreased expression of *vit-2* (Fig 4H, I) though other transcripts, including *act-1* and *ama-2* (Fig 4J,K), were unaffected by genotype.

We used WormCat (*19*, *20*) to identify functional classes within the transcripts that changed in *sams-1(lof)* animals with age, focusing initially on down-regulated gene products (Fig 5A, B). In these data, some categories of genes, such as mRNA function and CELL CYCLE, were enriched in both D1 and D7 (Fig 5A). This result is consistent with there being a common set of genes that decline with age in both WT and *sams-1(lof)* animals age. Since these changes occurred independently of *sams-1* we did not further investigate these categories. We focused instead on categories that were uniquely enriched in the D7 data, where there were particularly noticeable changes in METABOLISM (Fig 5A). This category includes many nuclear-encoded mitochondrial genes, which is reflected in the enrichment of “mitochondrial” genes in the METABOLISM category. Inspection of the individual gene products in the METABOLISM:Mitochondria set show broad downregulation of mitochondrial genes with age in *sams-1(lof)* animals. This downregulation was most pronounced in the mitochondrial ribosome (Fig 5B-F and Table S2). While some genes encoding ETC (electron transport chain) components were downregulated other pathways such as the citric acid cycle or ubiquinone synthesis were largely unaffected (Table S2). We also found enrichment of genes included in the Gene Ontology groups GO:00005739 (Mitochondrion) (Fig S3), which contains a broader array of genes encoding core mitochondrial-function proteins than the WormCat METABOLISM:Mitochondria set (*19*, *20*). These GO sets included genes such as tRNAs (Table S2) which could also be important for mitochondrial protein synthesis. In aggregate these data show that *sams-1* is required to maintain expression of mitochondrial genes as animals age.

**Figure 5:**
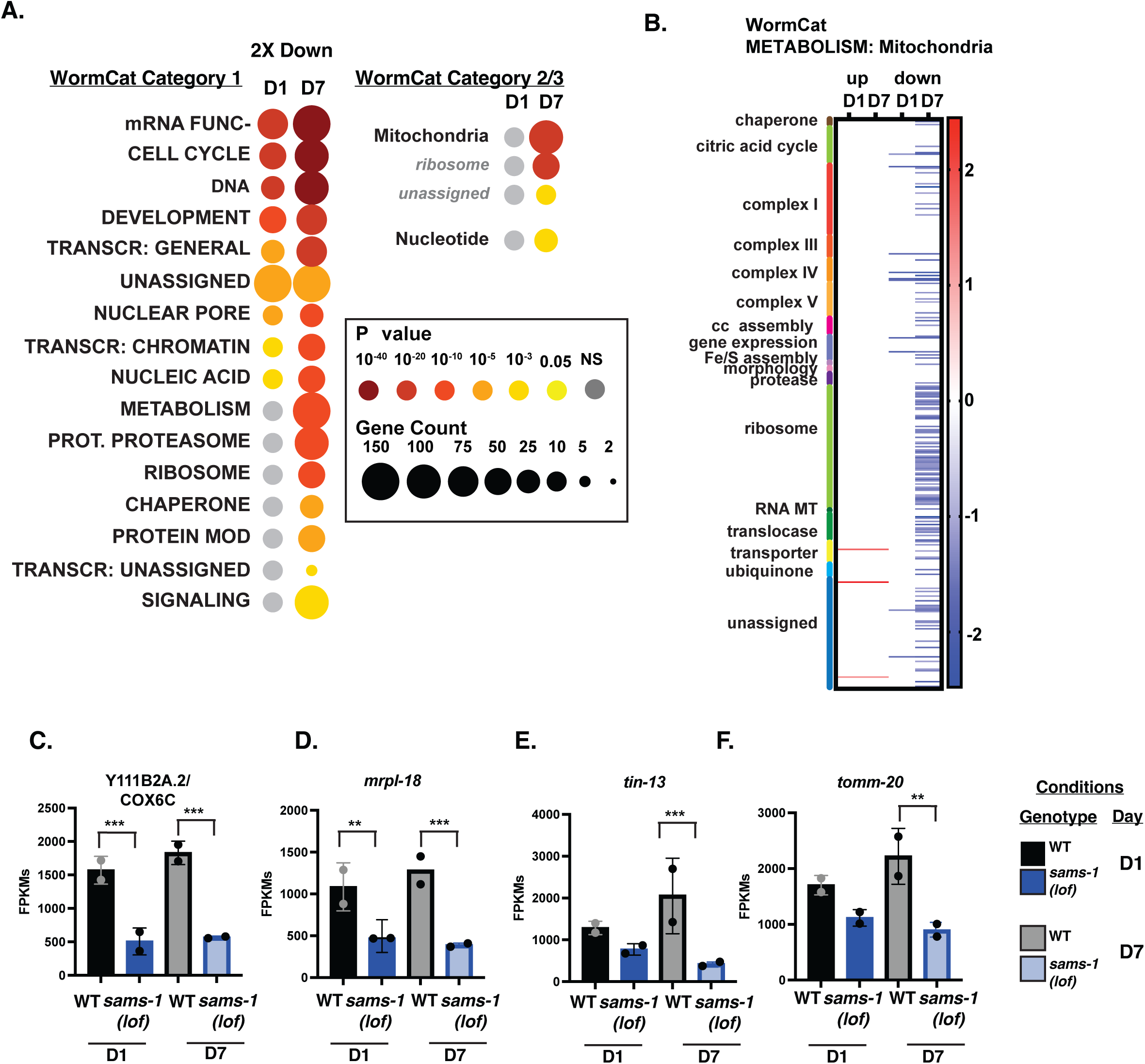
Genes for mitochondrial-targeted proteins are down regulated in aged populations of *sams-1(lof)* animals. **A.** WormCat pathway category analysis of 2 fold downregulated genes in *sams-1(lof)* animals at D1 and D7 of adulthood. Heat maps showing gene expression patterns for all genes in the WormCat category (**B**) METABOLISM: Mitochondria with category 3 divisions are noted on the left axis (**C**). Box and whisker plots showing FPKMs from D1 and D7 animals for nuclear encoded mitochondrial genes (**C-F**). Whiskers encompass standard deviation and the padjust value calculated by Deseq2 shows significance including a false discovery rate with * p <0.01, ** p <0.005, *** p 0.001.

### Limited overlap of *sams-1* upregulated genes with mitoUPR or autophagy regulators

WormCat analysis of genes with increased expression in early and aging *sams-1* animals showed multiple categories in common. For example, we found that gene products in the METABOLISM:lipid category were enriched among genes upregulated in *sams-1(lof)* animals (Fig 6A). This result is consistent with our earlier studies showing increased lipogenesis when *sams-1* is depleted (*7*). Similarly, the stress response category included many pathogen-related gene products, which we previously found were upregulated in D1 *sams-1(lof)* animals (*3*). Both lipid metabolism and pathogen stress response categories were more enriched in the set of downregulated genes in D7 *sams-1(RNAi)* animals, showing that these perturbations persist throughout the lifespan. We also noted some categories of genes that were only enriched in aged D7 *sams-1(RNAi)* animals: neuronal function, including subcategories enriched in synaptic function and neuropeptides, and transcription factors, which included several nuclear hormone receptors (*nhrs*).

**Figure 6:**
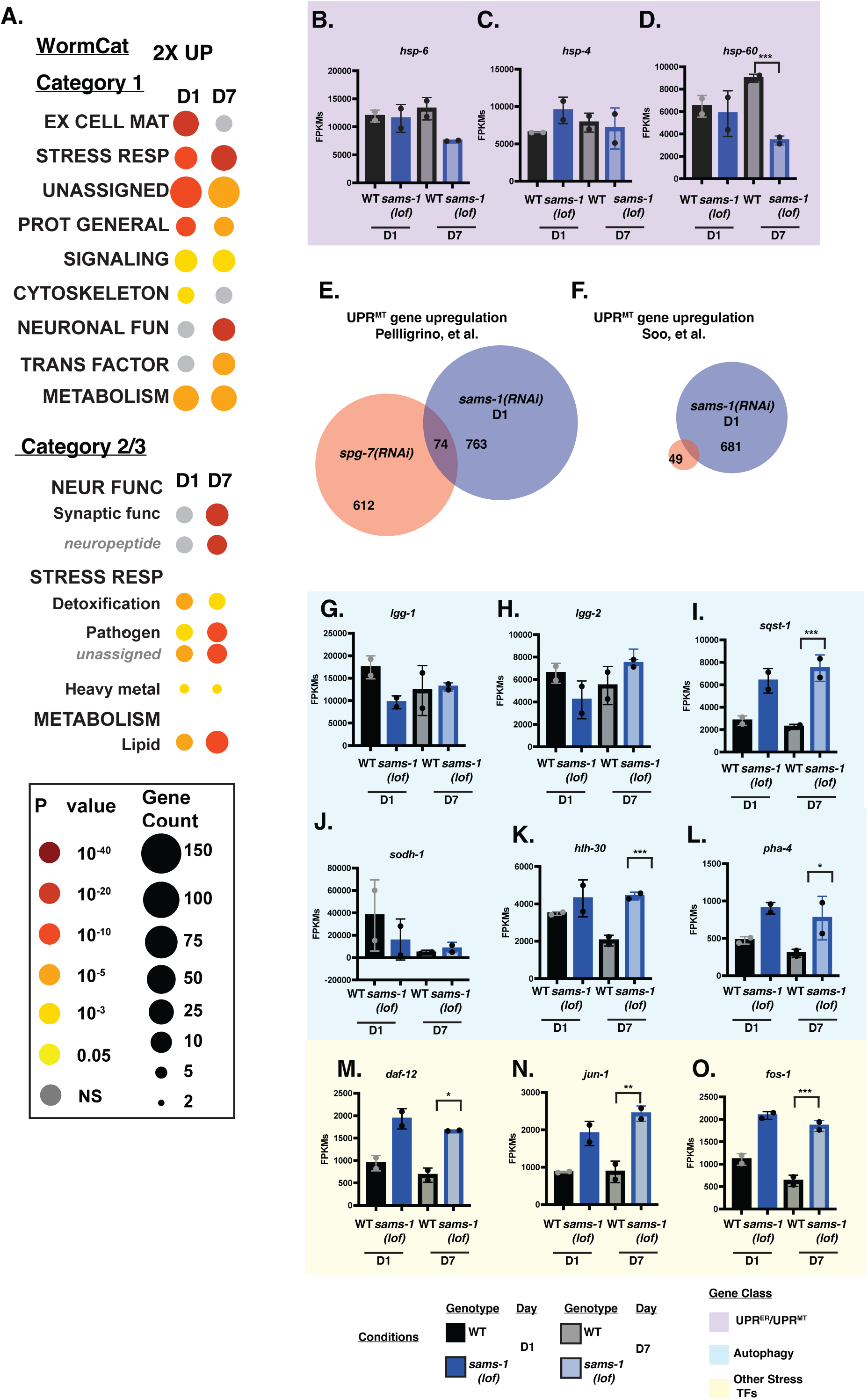
Limited evidence for upregulation of the UPR^Mito^ in young and aged *sams-1* animals. **A.** WormCat pathway category analysis of 2 fold up genes. Box and whisker plots showing FPKMs from D1 and D7 animals for UPR^Mito^ genes (purple box: **B-D**). Venn diagrams comparing sams-1 D1 upregulated genes with mitoUPR gene set in Pellegrino et al (**E**) or Soo, et al (**F).** Column graphs of FPKMs from RNAseq for autogphagy related genes (blue: **G-L**), other stress related transcription factors (yellow: **M-O**). Whiskers encompass standard deviation and the p-adjust value calculated by Deseq2 shows significance including a false discovery rate with * p <0.01, ** p <0.005, *** p 0.001.

Despite the downregulation of mitochondrial genes and lower levels of some metabolites (Fig 3, 5), we did not observe activation of the mito-UPR (mitochondrial unfolded protein response). The mito-UPR has been associated with increased lifespan and alterations in mitochondrial morphology (*21*). Several labs have reported increases in the expression of *hsp-6::gfp*, a transcriptional reporter for mito-UPR activation, in *sams-1(RNAi)* animals (*22–24*). Recently, a mitochondrial transporter SLC-25A26 has been suggested as a potential SAM transporter to the mitochondria, and its loss increased *hsp-6*::GFP reporter expression (*23*). However, there was no change in the abundance of *hsp-6* or *hsp-60* mRNAs in *sams-1(lof)* at D1 of adulthood in our RNAseq data (Fig 6 B-D), which is consistent with our previous microarray (*3*) and RNAseq (*4*) experiments. In fact, we found that these transcripts decrease rather than increase as animals age. More broadly, we found no significant overlap between gene products that change in our data set and multiple other studies where mito-UPR is activated (Fig 6E, F; (*25*, *26*)).

The discrepancy between our results and previous experiments using *hsp-6::gfp* could be a result of comparing the endogenous transcript with a high-copy reporter or could reflect persistence of GFP in those prior experiments from activation of the mitoUPR at an earlier point in development. In addition, the *hsp-*6::GFP strain has been shown to have increased stress sensitivity (*27*), suggesting that this multi-copy transgene may have neomorphic effects. We have noted a similar phenomenon with *hsp-4*, which is expressed upon activation of the ER UPR (*28*): although expression from the multicopy *hsp-4::gfp* reporter increases when *sams-1* is depleted, the abundance of endogenous *hsp-4* transcript is the same in *sams-1(lof)* and wild-type animals (Fig 6C; (*3*, *4*)). We therefore conclude, based on measurement of endogenous transcripts from multiple genes in this stress response, that the mito-UPR is not activated in *sams-1(lof)* animals and is therefore unlikely to contribute to observed lifespan and stress resistance phenotypes.

It has been suggested that increased autophagy in *sams-1(lof)* animals could contribute to increased lifespan and thermotolerance. Lim *et al.* showed that *sams-1(ok2946)* animals had increased expression of several autophagy-linked genes by qRT-PCR (*29*). However, this result is inconsistent with multiple independent RNAseq and microarray experiments, in which the expression of these autophagy genes did not change in *sams-1(lof)* or RNAi animals (*3*, *4*). Our data are consistent with these latter studies, as we did not observe an enrichment of genes in the WormCat AUTOPHAGY category among genes upregulated in *sams-1(lof)* animals (Fig 6A) and there were few changes in the expression of individual autophagy genes as animals aged (Fig 6G-L). The expression of autophagy regulators *pha-4* or *hlh-30* was similar at D1 in *sams-1(RNAi)* and wild-type animals, although these genes were upregulated in D7 *sams-1(RNAi)* animals (Fig 6 K-L). Furthermore, we found no significant differences in the expression of *lgg-1, lgg-2* or other effectors in *sams-1(lof)* animals, except for the cargo adaptor for selective autophagy, *sqst-1* (Fig 6 G-I). These results argue against a transcriptional induction of autophagy in *sams-1(lof)* animals. Further supporting this conclusion, there was no significant enrichment of H3K4me3 at the promoters of autophagy genes in genome-wide CUT&TAG in *sams-1(lof)* animals, excepting *sqst-1* (*5*, *30*). Although we did identify peaks of H3K4me3 in the same promoter regions used in prior ChIP-PCR experiments (*29*), these peaks were the same in *sams-1(lof)* and wild-type animals (Fig S4 A-G).

However, we did note that other genes included in STRESS RESPONSE categories was enriched in upregulated genes in D1 *sams-1(lof)* animals, which corroborates previous reports (*3*, *19*). We also found that aged D7 animals exhibited retention or enhancement in these categories (Fig 6A) and that multiple other stress-related transcription factors, including the AP1 components *fos-1* and *jun-1* and nuclear hormone receptor *daf-12*, increased as *sams-1* animals age, suggesting *sams-1* animals show broader increase along multiple stress-related categories as they age.

### Insufficient production of phosphatidylcholine leads to changes in mitochondrial morphology

Previous reports have linked changes in *sams-1* to mitochondrial fragmentation (*24*), which was proposed to result from depletion of SAM important for epigenetic regulation. Our finding of decreased mitochondrial metabolites and broad downregulation of mitochondrial genes in *sams-1(lof)* animals motivated us to visualize the mitochondrial networks as these animals aged. We found that the distribution of TOMM-20::mKate, a mitochondrially-localized fluorescent reporter (*31*) expressed in the intestine, was clearly disrupted in D1 *sams-1(RNAi)* animals. Instead of the wild-type reticular network, we observed many smaller mitochondrial bodies in *sams-1(RNAi)* animals (Fig 7B). This result is consistent with the fragmentation of body-wall muscle mitochondria observed in *sams-1(lof)* animals (*24*). In older control D7 animals, we observed mitochondrial fragmentation, consistent with previous observations (*32*). We also noted that *sams-1* animals showed a marked decrease in TOMM-20::mKate expression in both intestine and muscle tissues, suggesting an overall decrease in miotochondrial mass (Fig 7B, Fig S5A).

**Figure 7:**
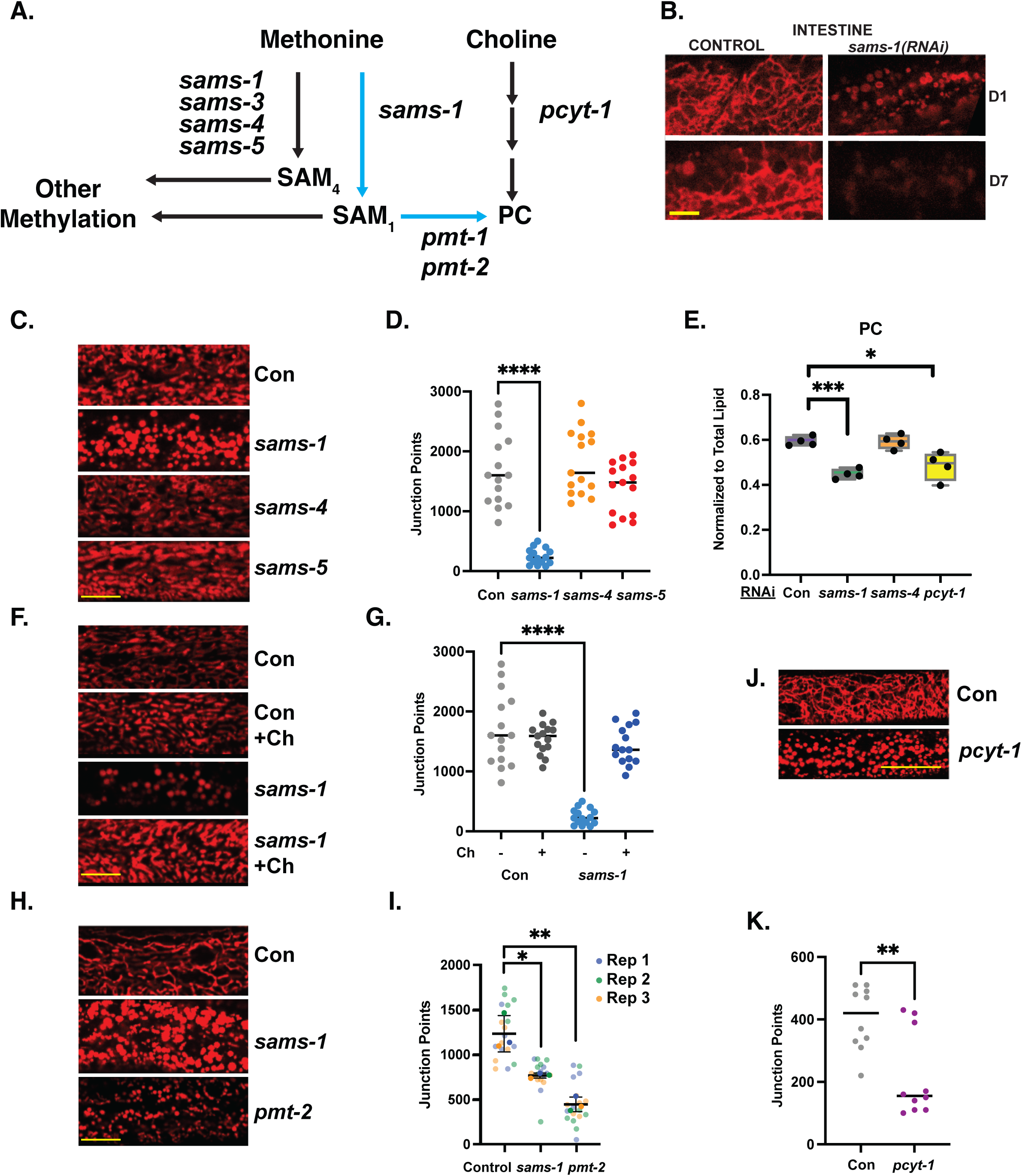
Changes in mitochondrial morphology in *sams-1* animals are linked to low PC. **A**. Schematic showing links between SAM and PC metabolism. **B.** *sams-1(RNAi)* causes changes in mitochondrial morphology at D1 of adulthood and loss of intestinal TOMM-20::mKate by D7. **C**. TMRE staining of D1 adult *C. elegans* after RNAi of each SAM synthase shows that mitochondrial effects are specific to *sams-1*, quantitation is in **D**. **E**. LCMS analysis shows that PC synthesis defects in low SAM are specific to *sams-1* (See also Figure S5, Table S3). **F.** Rescue of PC synthesis with dietary choline restores mitochondrial networks in D1 *sams-1(RNAi)* animals; quantitation is in **G**. **H.** RNAi of the PC methyltransferase *pmt-2* induces mitochondrial fission, visualized by TMRE staining, similar to reduction after *sams-1* RNAi. Quantitation is in **I. J.** *pcyt-1* RNAi TMRE quantitation is in **K.** Significance calculated by the Mann-Whitney test is shown with * p <0.01, ** p <0.005, *** p 0.001. Scale bar is 10 microns.

We observed similar mitochondrial fragmentation in *sams-1(RNai)* animals by staining with TRME, which accumulates in mitochondria (Fig 7C, D). Staining with TMRE also revealed that fragmentation of mitochondria is specific to *sams-1(lof)*, as we observed normal reticular mitochondrial morphology after RNAi of other SAMS enzymes *sams-3, 4* or *sams-5* (Fig 7C, D). One important function of SAMS-1 is to provide methyl groups required to maintain production of phosphatidylcholine (PC), an abundant membrane phospholipid (Fig 7A, (*33*)). Consistent with our previous studies, we measured a significant decrease in the amount of phosphatidylcholine in *sams-1(RNAi)* animals (Fig 7E;(*3*, *7*)). In contrast, *sams-4(RNAi)* animals have wild-type levels of phosphatidylcholine, indicating that SAM_4_ is not an efficient substrate for the PMT-1/PMT-2 phosphoethanolamine methyltrasferases. Furthermore, *sams-4* animals do not show changes in TG, DG or PG (Fig S5B-D) previously noted in *sams-1* animals (*34*, *35*) or exhibit any other significant changes in lipid class levels (Table S3). We conclude that the major SAM-dependent remodeling of lipid metabolism is specific to *sams-1*.

The deficit in phosphatidylcholine production and associated growth defect in *sams-1(lof)* animals can be rescued by dietary supplementation with choline (*3*, *7*, *36*, *37*), which can be incorporated into phospholipids through the Kennedy pathway (*33*). We found that supplementary choline rescued mitochondrial fragmentation in *sams-1(RNAi)* animals (Fig 7 F,G), suggesting that phosphatidylcholine is required to maintain a reticular mitochondrial network. To demonstrate that PC produced by SAM-independent pathways also produced the same phenotypes, we visualized the mitochondrial network in *pcyt-1(RNAi)* animals. PCYT-1, a choline-phosphate cytidylyltransferase, is the rate-limiting enzyme in the *de novo* (Kennedy) pathway, which uses choline and thus does not depend on methylation (*33*). PC levels were similar after *sams-1* or *pcyt-1(RNAi)* (Fig 7E), suggesting these pathways contribute broadly to PC production in *C. elegans*. TMRE staining revealed a fragmented mitochondrial network in *pcyt-1(RNAi)*, indistinguishable from our observations in *sams-1(RNAi)* animals (Fig 7J, K). Further supporting this notion, we found that knockdown of the phosphoethanolamine methyltransferase *pmt-2* also increased mitochondrial fragmentation (Fig 7H, I). Notably, *sams-4* was dispensable for PC production (Fig7E) and did not display mitochondrial fragmentation (Fig7C), suggesting the specific role for SAMS-1 in PC production is the mechanistic key to this phenotype. Our results strongly support the model that differences in phospholipids underlie the mitochondrial fragmentation in these animals.

We next examined levels of other lipids that may have significant impacts on mitochondrial function (*38*). In our lipidomics assays, we noted that PE (phosphatidylethanolamine) and cardiolipin (CL) levels were similar in control and *sams-1* RNAi animals (Fig S5C, H; Table S3), although recovery of CL was limited.

Nevertheless, this result suggests that these lipids did not contribute to the mitochondrial fragmentation we observed. We did note an increase in sphingosines (SPH) along with lipid-conjugated metabolites with important links to mitochondrial functions such as Acyl carnitines and Coenzyme Q (ubiquinones) after RNAi of either *sams-1* or *pcyt-1* (Fig S5G, I, J). One possibility is that these changes reflect a compensatory program in these altered mitochondria, though we cannot exclude direct roles for these lipids at this time.

### Fragmented mitochondria in sams-1 animals are targeted to autophagosomes

Morphological dynamics of mitochondria are tightly linked to stress responses; for example, in *C. elegans* mitochondrial dynamics can affect mitophagy and heat shock survival (*39*). In *C. elegans* larvae exposed to heat stress, mitochondrial fragmentation protects animals from heat stress and requires the dynamin ortholog *drp-1* (*39*). In contrast, vitamin B12 deficiency causes mitochondrial fragmentation in *drp-1(tm1108)* mutant animals (*24*), suggesting that fragmentation linked to 1CC function does not require DRP-1 activity. Mitochondrial fusion, on the other hand, is driven by *fzo-1*, a mitochondrial membrane localized GTPase. We confirmed that in our experiments *drp-1(RNAi)* animals were more sensitive to heat stress and that, conversely, *fzo-1(RNAi)* animals were resistant (Fig 8A). We next used RNAi to deplete *drp-1* or *fzo-1* in *sams-1(lof)* animals. We found that, similar to the wild-type, *sams-1(lof);drp-1(RNAi)* animals were more sensitive to heat stress than *sams-1(lof)* alone (Fig 8B). The thermal tolerance of the *sams-1(lof)* animals did not change when *fzo-1* was depleted (Fig 8B), in contrast to increased survival in wild-type (Fig 8A), perhaps because the mitochondria were fragmented even without loss of *fzo-1*.

**Figure 8:**
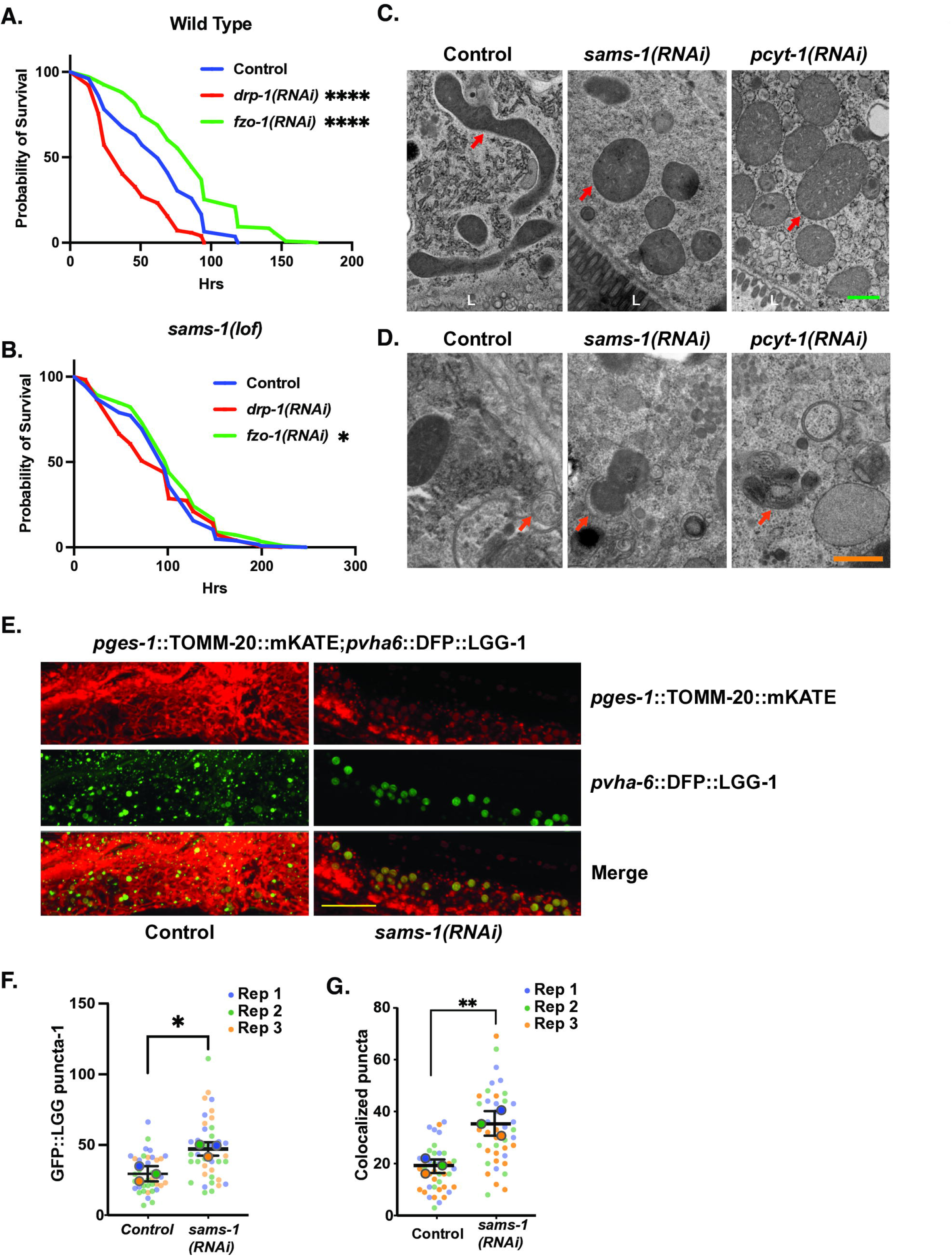
Increased mitophagy in *sams-1(RNAi)* animals links changes in PC to heat shock survival. Average of three survival assays comparing effects of heat shock on wild type (**A**) or *sams-1(lof)* (**B**) animals undergoing RNAi for *drp-1* or *fzo-1*. Panel A confirms results from Chen (2021). TEM images of Control, *sams-1* or *pcyt-1* RNAi animals comparing mitochondrial morphology, localization and localization within intestinal cells (**C**) and showing examples of mitochondria in proximity to double membrane autophagosome-like structures (**D**). Scale bar is 10 microns**. E.** Confocal projections obtained on a Nikon Spinning disc of intestines from *sams-1(RNAi)* animals showing increased numbers of LGG-1::GFP autophagosome puncta and overlap with TOMM-20::mKate marked mitochondria. Scale bar is 500 nm. Quantification of GFP::LGG-1 puncta is in **F** and of overlap between GFP::LGG-1 puncta and TOMM-20::mKate in **G**. Significance calculated by the Mann-Whitney test is shown with * p <0.01, ** p <0.005, *** p 0.001.

Mitochondrial fragmentation is a key step to initiate the specific degradation of mitochondrial components by autophagy, or mitophagy (*40*). We used high pressure freeze fixation/transmission electron microscopy (TEM) to examine the ultrastructural characteristics of mitochondria after *sams-1(RNAi)*. Whereas intestinal mitochondria in control animals where elongated with visible cristae and were dispersed throughout the cytoplasm, we noted that the mitochondria of *sams-1(RNAi)* animals were preferentially distributed along the intestinal lumen, had a rounded or bloated morphology, and barely discernible internal morphology (Fig8C, FigS6A). We observed a similar effect in *pcyt-1(RNAi)* animals, further supporting our assertion that *sams-1* mitochondrial phenotypes are a result of low PC. PC is a major constituent of other cellular membranes, and we noted tubulation of ER membranes in addition to the mitochondrial effects. These fragmented mitochondria in *sams-1(RNAi)* animals were often observed in close proximity to cupped-shaped double membrane structures, suggesting that mitophagy could be increased. This is consistent with genetic studies that showed the autophagy regulator *bec-1* is required for full lifespan extension in *sams-1(ok2946)* animals and that these animals showed increased LGG-1::GFP puncta increased in body wall muscle and other tissues (*29*).

In order to quantitatively measure if autophagic processes were linked to the changes in mitochondrial morphology, we knocked down *sams-1* in a strain where intestinal autophagosomes were marked with DFP::LGG-1 (*41*) and mitochondria with TOMM-20::mKate (*41*). In our *ges-1*::TOMM-20mKate micrographs, mitochondria in *sams-1(RNAi)* animals clustered near the basal side of the intestinal cells as in our TEM. In addition, lighter, less refractory puncta were present closer to the apical side (Fig 8E, FigS6A). We therefore hypothesized that mitophagy could be occurring in these animals to target the fragmented mitochondria. Similar to Lim et al. (*29*) we noted increased large LGG-1::GFP puncta in *sams-1(RNAi)* animals (Fig 8E, F). Many of the LGG-1::GFP puncta colocalized with mitochondria in *sams-1(RNAi)* animals (Fig 8E, G).

Mitochondria associated with LGG-1::GFP appeared more spherical or bloated and stained less intensely with TOMM-20::mKate than other mitochondria in the cell (Fig8E). These results suggest that in *sams-1(lof)* animals, inadequate production of PC leads to mitochondria fragmentation and subsequent degradation by mitophagy (see model, SFig6B). This mitochondrial fragmentation phenotype is similar to reduction in PCs through loss of the PC transfer protein STARD7, which causes mitochondrial disfunction associated with a seizure disorder in humans (*42*). Loss or reduction of STARD7 alters mitochondrial morphology with increased blebbing and loss of cristae structure (*42*, *43*). Moreover, the mitochondrial morphology after loss of STARD7 is also independent of dynamin-related GTPases (*43*) and increased mitophagic flux in C1C12 muscle cells (*44*). Taken together, our results suggest that that the mitochondrial fragmentation in *sams-1* animals is upstream of the cytoskeletal mitochondrial remodeling pathways, and that mitochondrial fragmented through changes in membrane lipid may be primed to enter mitophagy pathways.

## Discussion

We have previously found that SAM synthases have distinct effects on lifespan, heat stress survival, gene expression profiles and H3K4me3 patterns (*5*). Here, we find that SAMS-1 contributes broadly to metabolism, including a mitochondrial β-oxidation-like pathway and in synthesis of the phospholipid PC. In contrast to SAM_1_, our studies indicate that SAM_4_ may play a more restricted role, perhaps serving primarily as a substrate for a more limited set of histone or other protein methylation reactions. These results support the idea that SAM generated by different SAMS enzymes is used for specific cellular and metabolic processes. Taken together, our study demonstrates that effects of lowering SAM or impacting 1CC metabolism can act on multiple cellular pathways to alter aging or stress responses. Our study points to a need to integrate multiple methylation-dependent mechanisms and the potential for synthase-specific effects when determining how low SAM may exert phenotypic effects.

Long-lived *sams-1* animals show loss of mitochondrial gene expression as they age, along with mitochondrial fission and loss of mitochondria through mitophagy, which we demonstrate is mechanistically linked to PC. This is reminiscent of mammalian studies showing loss of MAT1A in mammalian liver impacts mitochondria and may underlie changes occurring in alcoholic fatty liver disease (*55*). We suggest that the SAM/PC axis is key to effects on mitochondria which link heat stress survival and aging in these animals. In mammals, mutations in choline kinase controlling initial steps in PC synthesis are associated with changes in mitochondrial morphology and mitophagy in mouse muscle tissue (*50*) and human disease (Congenital muscular dystrophy with mitochondrial structural abnormalities, CMDmt), underscoring the importance of understanding these links between metabolism, mitophagy and stress resistance/lifespan.

### SAMS-1 dependent metabolites impact mitochondrial morphology

SAM levels could impact metabolic pathways as a substrate, allosteric regulator, or by influencing chromatin modification and gene expression. In our previous work (*5*), we observed that genes in the propionate shunt, a β-oxidation-like pathway, lost H3K4me3 and heat-shock associated gene expression. These results extend these findings, showing that multiple metabolites in this pathway are impacted.

There are clear links between the regeneration of methionine and SAM synthesis though vitamin B12 and the detoxification of hydroxy propionate (3HPP) (*56–59*). Recent studies have linked increased 3HPP with defects in mitochondrial morphology including sheet like structures and vacuoles (*51*). These can be rescued by B12 or the low branched chain amino acid (BCAA) fatty acid diet of the *E. coli* strain used for *C. elegans* RNAi studies. While B12 and BCAA metabolism can affect the 1CC, *sams-1(RNAi)* animals showed no significant differences in the BCAAs leucine or isoleucine, although valine levels dropped. This suggests that other mechanisms, such as the reductions in levels of H3K4me3 and expression in genes for metabolic enzymes in this pathway (*5*) accounts for the observed decreases in 3HPP.

We have demonstrated that reduced PC production in *sams-1* animals is closely associated with mitochondrial fragmentation and increased mitophagy. PC forms a major part of most cellular membranes, so the mitochondrial effects we observe could arise from its role as a metabolic precursor of other complex lipids or by altering membrane dynamics/membrane protein function. PC is used in the synthesis of multiple other lipids including cardiolipin, which plays an important role in mitochondrial membranes. However, we found that CL levels are maintained in *sams-1* animals, suggesting that perturbation of CL function do not lead to the mitochondrial fragmentation we observe in *sams-1(RNAi)* animals. However, we did note several changes in the abundance of other lipids linked to mitochondrial function such as PG, PI, SPH, CoQ and Acyl Carnitines when SAMS-1 is depleted. These lipid classes also increased in *pcyt-1* RNAi animals. It may be that TG, DG, PG, SPH and CoQ lipids accumulate when PC is limited due to compensatory changes to rebalance mitochondrial function. PC has also been linked to functional changes in mitochondria in other contexts. Recent studies found that the CoQ variants compete with PC for binding the lipid transfer protein STARD7 (*54*). Finally, using a screen in yeast for regulators of autophagy/mitophagy induced by low methioninem= Sutter and Tu (*67*) identified ChoP (cholinephosphotransferase), highlighting deep links of PC production to the autophagy/mitophagy machinery.

Limitations in available PC may also affect mitochondrial morphology and membrane potential through effects on membrane dynamics (*54*, *55*). PC plays a critical role in membrane stability and curvature, as its cylindrical structure promotes bilayer formation (*56*). The effect of decreased in PC on membrane dynamics are well studied in lipid droplet biology (*57*), as PC depleted membranes are less amenable to curvature, with larger and more regular formation of lipid droplets. This is consistent with the larger, less networked mitochondria in both the *sams-1* and *pcyt-1* RNAi animals. In mitochondria, low PC may also favor fission rather than maintenance of complex fused mitochondrial networks and alter modifications of mitophagy regulators such as Atg8 (*58*) or Atg44 (*59*), which could influence incorporation of damaged mitochondria into the autophagy machinery.

We previously showed that in *C. elegans* and mammalian cells, low PC blocked ARF-1 cycling (*35*), which can affect multiple membrane organelles. Studies from the Sprang lab make a strong case for the importance of ARF-1/ Arf1/ARF1 in mitochondrial dynamics in *C. elegans*, yeast and mammalian cells (*60*, *61*). However, unlike our observations in *sams-1(lof)* animals, loss of *arf-1* does not promote mitochondrial fission. This result suggests that ARF-1 is not the key effector of mitochondrial morphology when PC is reduced. Effects of low PC are likely to affect organelles differently, as the ratios to PC to other lipids vary, with highest levels in the ER, decreases in Golgi, mitochondria and lysosomes and lowest in the plasma membrane (*62*). Thus, SAM synthase specific effects on PC levels appears to be the mechanistic key to *sams-1* effects on mitochondrial morphology. Our study highlights the importance of demonstrating downstream roles for SAM and suggest that links to specific methylation targets are essential to understand its importance in lifespan and stress responses.

## Materials and Methods

### *C. elegans* strain and culture, RNAi and stress applications

The N2 strain or *sams-1(ok3033)* (*Caenorhabditis* Genetics Center) was used for all experiments. Culture conditions are as previously described (*5*). For the metabolomics and heat shock experiments, adults were bleached onto RNAi plates and allowed to grow to young adults at 15°C before heat shock. Animals were collected in S-Basal, washed and frozen at –80°C until processing. For all other experiments, animals were maintained at 20°C. For heat shock survival assays, animals were bleached onto RNAi plates and allowed to grow to young adults at 15°C. Approximately 25-30 animals were moved onto 35mm plates in triplicate, resulting in 75-90 animals in each biological replicate. These plates were placed in a 37°C incubator for 2 hours, then moved to 20°C for the remainder of the assay. Starting the next day, each plate was checked for dead animals by looking for movement and, failing that, gentle prodding of the worm. Dead worms were removed and scored. Animals that died from internally hatched embryos (bagging) or died desiccation were excluded from the final scoring. Three independent non blinded biological replicates were carried out and Kaplan-Meir curves were generated with GraphPad Prism v8.0

## Relative targeted metabolite profiling

### Sample Preparation

Aqueous metabolites for targeted liquid chromatography–mass spectrometry (LC-MS) profiling of 80 fly samples were extracted using previously described protein precipitation method (*63–65*). Briefly, samples were homogenized in 200 µL purified deionized water at 4 °C, and then 800 µL of cold methanol containing 124 µM 6C13-glucose and 25.9 µM 2C13-glutamate was added. Internal reference standards were added to the samples to monitor sample preparation. Next, samples were vortexed, incubated for 30 min at-20 °C, sonicated in an ice bath for 10 min, centrifuged for 15 min at 18,000 x g at 4 °C, and then 600 µL of supernatant was collected from each sample (protein precipitate was used to do BCA assay for data normalization). Lastly, recovered supernatants were dried on a SpeedVac and reconstituted in 0.5 mL of LC-matching solvent containing 17.8 µM 2C13-tyrosine and 39.2 3C13-lactate, and internal reference standards were added to the reconstituting solvent to monitor LC-MS performance. Samples were transferred into LC vials and placed in a temperature-controlled autosampler for LC-MS analysis.

### LC-MS Assay

Targeted LC-MS metabolite analysis was performed on a duplex-LC-MS system composed of two Shimadzu UPLC pumps, CTC Analytics PAL HTC-xt temperature-controlled auto-sampler and AB Sciex 6500+ Triple Quadrupole MS equipped with ESI ionization source (*65*). UPLC pumps were connected to the autosampler in parallel and were able to perform two chromatography separations independently from each other. Each sample was injected twice on two identical analytical columns (Waters XBridge BEH Amide XP) performing separations in hydrophilic interaction liquid chromatography (HILIC) mode. While one column was performing separation for MS data acquisition in ESI+ ionization mode, the other column was equilibrated for sample injection, chromatography separation and MS data acquisition in ESI-mode. Each chromatography separation was 18 min (total analysis time per sample was 36 minutes), and MS data acquisition was performed in multiple-reaction-monitoring (MRM) mode. The LC-MS system was controlled using AB Sciex Analyst 1.6.3 software. The LC-MS assay targeted 361 metabolites and 4 spiked internal reference standards. Measured MS peaks were integrated using AB Sciex MultiQuant 3.0.3 software. Up to 201 metabolites and 4 spiked standards were measured across the study set, and over 90% of measured metabolites were measured across all the samples. In addition to the study samples, two sets of quality control (QC) samples were used to monitor the assay performance and data reproducibility. One QC [QC(I)] was a pooled human serum sample used to monitor system performance, and the other QC [QC(S)] consisted of pooled study samples, which were used to monitor data reproducibility. Each QC sample was injected once for every 10 study samples. The data were highly reproducible with a median coefficient of variation (CV) of 4.0 %.

## Metabolite Data Analysis

The raw metabolomics data was first preprocessed to ensure analysis of only high quality data (*66*). Metabolites that were not captured in 75% or more of the samples were excluded from the analysis. Next, metabolites solely from bacteria sources were identified and removed by comparing the mean of pooled blanks with the mean of pooled samples. Quality control was performed on a pooled sample where the same metabolites were repeatedly measured. If the coefficient of variation was > 20%, metabolites were excluded from downstream analysis (*66*). This dataset was uploaded onto the Metaboanalyst v5.XX web platform (*67*). Metaboanalyst to normalize our data before identifying changing metabolites with respect to each SAM synthase or after heat shock. Missing values were handled with the default setting (remove feature with >50% missing values and otherwise replace with 1/5 of the minimum positive value for each variable). Data was ultimately normalized by sum and log10 transformation after examining the distribution of each normalization combination (*68*). This data was downloaded and fed into GraphPad Prism v10.0.0 where a two-way ANOVA was performed. False discovery rate was controlled with the Two-stage step-up method of Benjamini, Krieger and Yekutieli and a desired false discovery rate of 0.05. Graphs were visualized in Prism except for the principal component analysis was performed in R v4.2.1.

### Lipidomics

*C. elegans* samples were extracted using a modified Folch procedure including Internal Standards (Splash Mix; Avanti Polar Lipids). Samples were resuspended based on sample weight. ***C-MS/MS*** Thermo Accucore C30 150mm analytical column was used in the positive and negative ionization modes to acquire data. Data were analyzed using LipidSearch 5.1. Background filtering was performed using solvent blanks (no extraction blanks submitted, which are the ideal sample to filter out background). Data were filtered within the software according to pre-determined parameters within the software. Lipid Class data was used to generate a total lipid signal, Data shown were normalized to total lipid signal. Statistical analysis and visualization were performed in Graphpad prism.

### RNA sequencing

RNA for deep sequencing was purified by Qiagen RNAeasy. Duplicate samples were sent for library construction and sequencing at BGI (China). Raw sequencing reads were processed using a Nextflow pipeline (https://github.com/DanHUMassMed/RNA-Seq-Nextflow). The raw read pairs were first aligned to *C. elegans* reference genome with ws245 annotation. The RSEM method was used to quantify the expression levels of genes and Deseq was used to produce differentially expressed gene sets with more than a twofold difference in gene expression, with replicates being within 0.05 in a Students T test and a False Discovery Rate (FDR) under 0.01. Statistics were calculated with DeBrowser (*69*). Venn Diagrams were constructed by BioVenn (*70*).WormCat analysis was performed using the website https://www.wormcat.com/ (*19*, *20*)and the whole genome annotation version 2 (v2) and indicated gene sets. PCA was conducted by using *prcomp* in R and graphed with *ggplot* in R studio. GEO accession numbers are pending.

### Microscopy TMRE staining

Plates with TMRE were prepared by dripping 10 μM TMRE onto the bacterial lawn and allowing the plate to dry in a hood immediately before use. Embryos from 20-25 animals were recovered by hypochlorite treatment on TMRE plates and imaged when the animals reached the young adult stage.

### Confocal imaging

*C. elegans* were imaged on a Leica SPE6 or Nikon Spinning Disc as noted with identical gain or exposure settings within each experiment. Images were quantitated in FIJI with the following pipelines: 1) TMRE staining with MitoMapr (*71*), 2) PunctaProcess (https://github.com/DanHUMassMed/puncta_process_plugin) for counting GFP::LGG-1 and 3) Colocalization of GFP::LGG-1 and TOMM-20::mKate with the Fiji Image calculator, followed by Punta Process to quantify the overlapping pixels. Graphs and statistical analysis were completed in Graphpad Prism.

### Transmission Electron Microscopy

A suspension of *C. elegans* (tetramizole/ M9) were placed in the 100 μm deep side of type A 6 mm Cu/Au carriers (Leica), sandwiched with the flat side of type B 6 mm Cu/Au carriers (Leica) and frozen in a high-pressure freezer (EM ICE, Leica). This was followed by Freeze Substitution (FS) in a Leica EM AFS2 unit cooled down to-90°C. For Freeze Substitution Procedure and Embedding, the sandwiches with frozen samples were transferred under LN2 into cryovials containing frozen FS media (Acetone containing 1% OsO4/1% glutaraldehyde/1% water). The vials are placed into the precooled AFS2 unit, the lids are screwed loosely onto the vials to permit safe evaporation of excess N2 gas and after about 1 h the lids are tightened, and FS is started. During the freeze substitution, the temperatures increased to - 90°C for 36-48 hrs to 60°C at a rate of 5°C per hour (6h). After-60°C for 6 hrs, samples were warmed up to-30°C at a rate of 5°C per hour (6h). Similarly, after 3 hrs at - 30°C samples were warmed up to 0°C at a rate of 5°C per hour over 6 hours. Finally, samples were warming up to 20°C at a rate of 5°C per hour. After reaching room temperature the samples were rinsed three times with acetone (10 min each), one time with propylene oxide (PO) and infiltrated in a 1:1 mixture of PO:TAAB Epon overnight. The next day, specimens were transferred to 100% TAAB Epon, incubated 2 h at RT then transferred to fresh TAAB Epon in an embedding mold or embedded flat between sheets of Aclar, plastic and moved to the oven to polymerize at 60°C for 48hr.

## Acknowledgements

We would like to thank Drs. Cole Haynes, Marian Walhout (UMASS) and Miriam Greenberg (Wayne State) for helpful discussion. We appreciate the assistance of Dr. Marie Bao, Life Science Editors, for help in preparing the manuscript, along with Dr. Nils Grotehans and members of the Walker lab for helpful comments. We thank the lab of Dr. Jin Zhang (UMASS) for use of the Nikon Spinning Disc for confocal microscopy and to the UMASS Metabolomics core for lipidomics. Metabolomics were also performed at the University of Washington Nathan Shock Center of Excellence in the Basic Biology of Aging and transmission electron microscopy at the Harvard Medical School microscopy core. Some strains were provided by the CGC, which is funded by NIH Office of Research Infrastructure Programs (P40 OD010440).

## Funding

National Institutes of Health grant R01AG0686701 (AKW)

National Institutes of Health grant 1R01AG053355 (AKW, DLM)

United States Department of Agriculture cooperative agreement USDA/ARS 58-8050-9-004 (DP)

National Institutes of Health grant University of Washington Nathan Shock Center of Excellence in the Basic Biology of Aging (NIH P30AG013280)

Searle Scholar’s Program (JBS)

AFAR Junior Faculty Award (JBS)

Smith Family Foundation (JBS)

## Supplemental Figures and Tables

**Figure S1: Schematic of metabolic pathways represented in data from targeted metabolomics comparing SAM synthase knockdown in basal and heat shocked *C. elegans*, based on WormPaths diagrams.** Pathways were adapted from WormPaths (*16*). Colored boxes correspond to metabolites shown in individual graphs. Bolded metabolites are represented in targeted metabolomics.

**Figure S2: Comparison of individual metabolite levels from targeted metabolomics**. Box and whisker plots showing individual metabolites from targeted metabolomics comparing heat shocked *sams-1 and sams-4(RNAi)* animals (**A-H**). Colored boxes show location of selected metabolites on **Figure S2**. Significance was determined by two-way repeated measures ANOVA. ns: q-value ≥ 0.05 *: q-value < 0.05, **: q-value < 0.01, ***: q-value <.001, ****: q-value > 0.0001. Color blocks map to areas on metabolic map (Figure S2, Figure 2 and Figure 3).

**Figure S3: Distribution of *sams-1(lof)* upregulated genes during aging in GO.** Heat map of GO: 00005739 (Mitochondrion) category.

**Figure S4:** Browser tracks showing H3K4me3 peaks from Godbole, et al. eLife (2023) of autophagy-related genes after *sams-1* and *sams-4(*RNAi) for *hlh-30* (**A**), *pha-4* (**B**), *sqst-1*(**C**), *lgg-1* (**D**), *lgg-2* (**E**), *atg-4.1* (**F)**, and *sodh-1*(**G**). Locations for primer sets used in Lim et al 2023 are boxed.

**Figure S5: Changes in mitochondrial morphology and lipid levels occur after loss of *sams-1*. A.** Confocal projection shows mitochondrial fission progression to loss from D1 to D7 in body wall muscle expressing TOMM-20::mKATE animals after *sams-1(RNAi)*. LCMS analysis comparing lipid class levels after *sams-1, sams-4* or *pcyt-1(RNAi)* for TG (triglycerides) (**B**), PE (phosphatidylethanolamines) (**C**), diglycerides (DG, **D**), phosphatidylglycerols (PG, **E**), Phosphatidylinstitols (PI, **F**), Acylcarnatines (AcCa, G), cardiolipins (CL, **H**), Ubiquinone isoforms (CoQ, **I**) and sphingolipids (SPH, **J**).

**Figure S6. Changes in mitochondrial morphology and cellular localization occur after loss of *sams-1*. A.** TEM images of Control, *sams-1* or *pcyt-1* RNAi animals comparing mitochondrial morphology, localization and localization within intestinal cells. Scale bar is 500 nm. **B**. Model.

**Supplemental Table 1: Targeted metabolomics of Control, *sams-1(RNAi)* and *sams-4(RNAi)* animals in basal and heat shock conditions.**

**Tab 1:** Metabolites detected by LCMS in targeted analysis. Each metabolite is cross referenced by HMB ID (Human Metabolome Database), common name, pubChem and KEGG databse IDs, and WormPaths (*16*) identification. Classyfire (*72*), RefMet (*73*) and WormPaths classifications are included where available

**Tab 2:** LogFold changes for each metabolite. Up (peach) or down (light blue) changes were considered significant if over two fold with a p value > 0.05.

**Supplemental Table 2: RNA seq comparing gene expression during aging in wild type and *sams-1(lof)* animals.** Tabs contain Deseq output, comparisons to published mitoUPR and GO: Mitochondrion gene list and comparisons of WormCat enrichment for categories 1, 2 and 3. **List of Tabs: Tab 1:** N2_sams1_D1_all, **Tab 2:** N2_sams1_D7_all**, Tab 3:** D1_D7_comp**, Tab 4**: GO-0005739**. Tab 5**: WormCat1**, Tab 6**: WormCat2**. Tab 7:** WormCat3**. Abbreviations**: NC: No change, NF: Not found, NS: Not significant. NV: No value.

**Supplemental Table 3: Lipid classes in Control, *sams-1, sams-4* and *pcyt-1* animals.** Lipid classes detected by LCMS and normalized to total lipid levels. Significance determined by paired Students T test.

