## Supplementary figures and images for "Functional Specialization of S-Adenosylmethionine Synthases Links Phosphatidylcholine to Mitochondrial Function and Stress Survival"

### Supplemental Figure 1

Figure S1
Munden, et al.

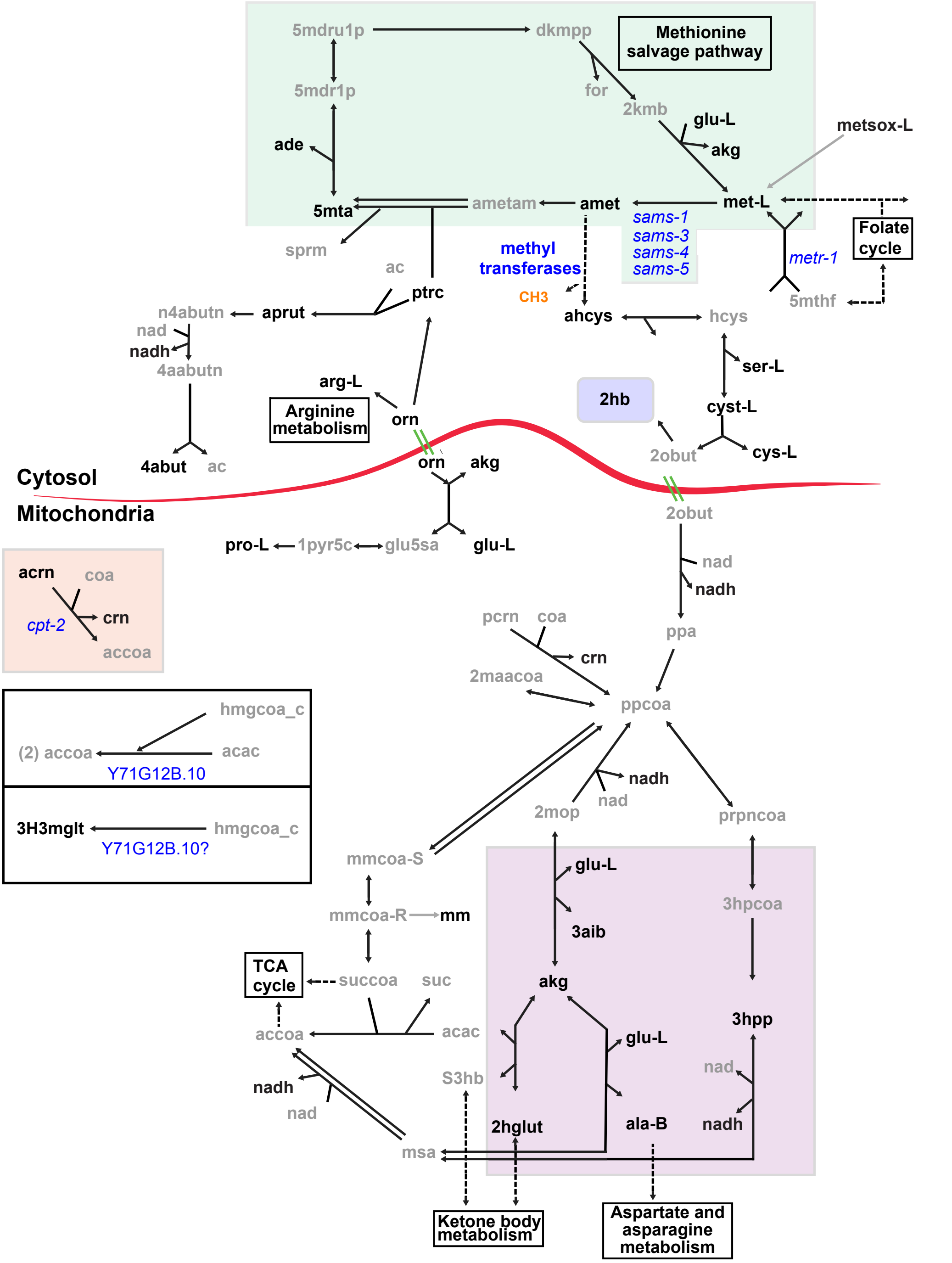

### Supplemental Figure 2

Fig S2
Munden, et al.

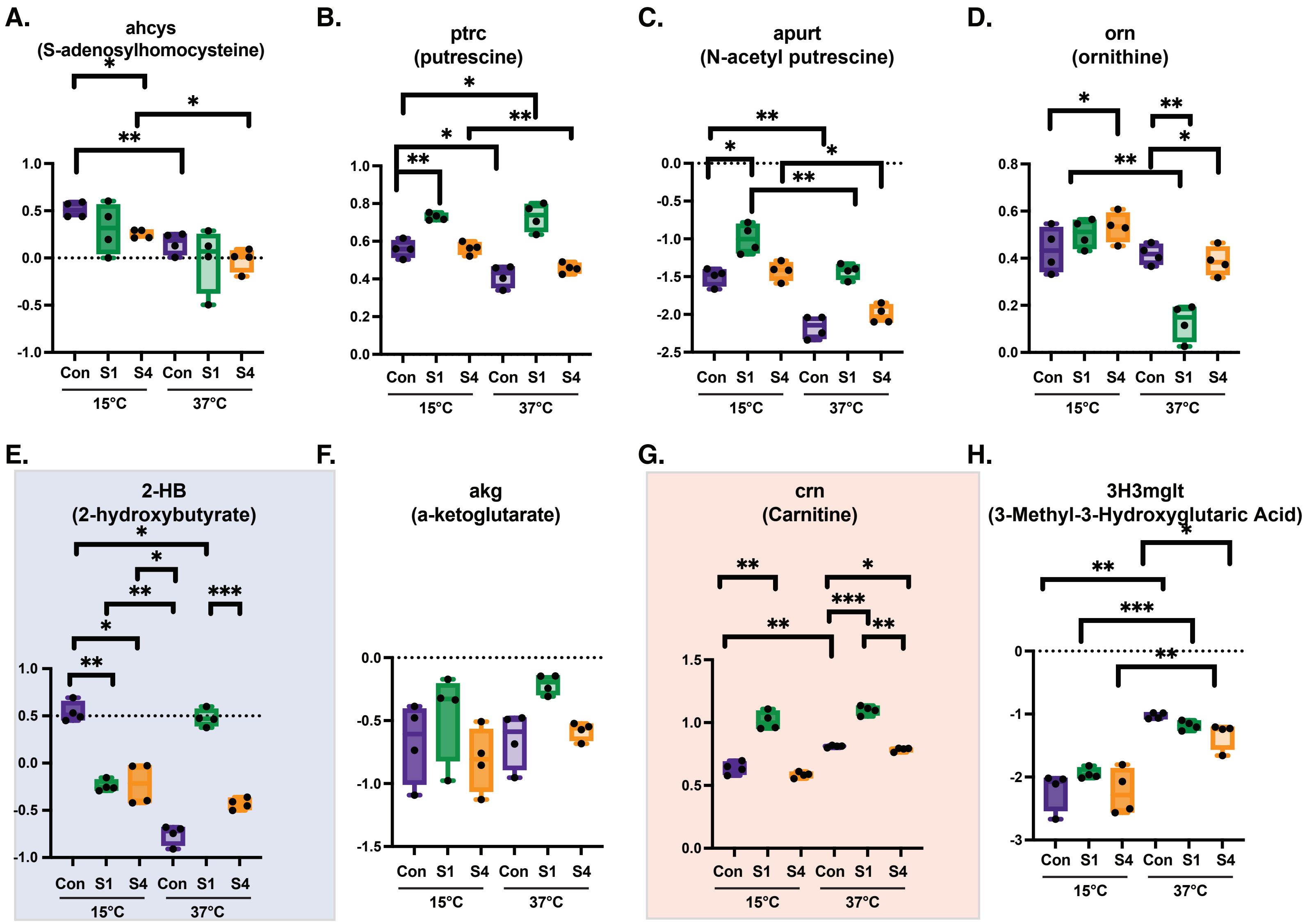

### Supplemental Figure 3

Fig S3
Munden, et al.

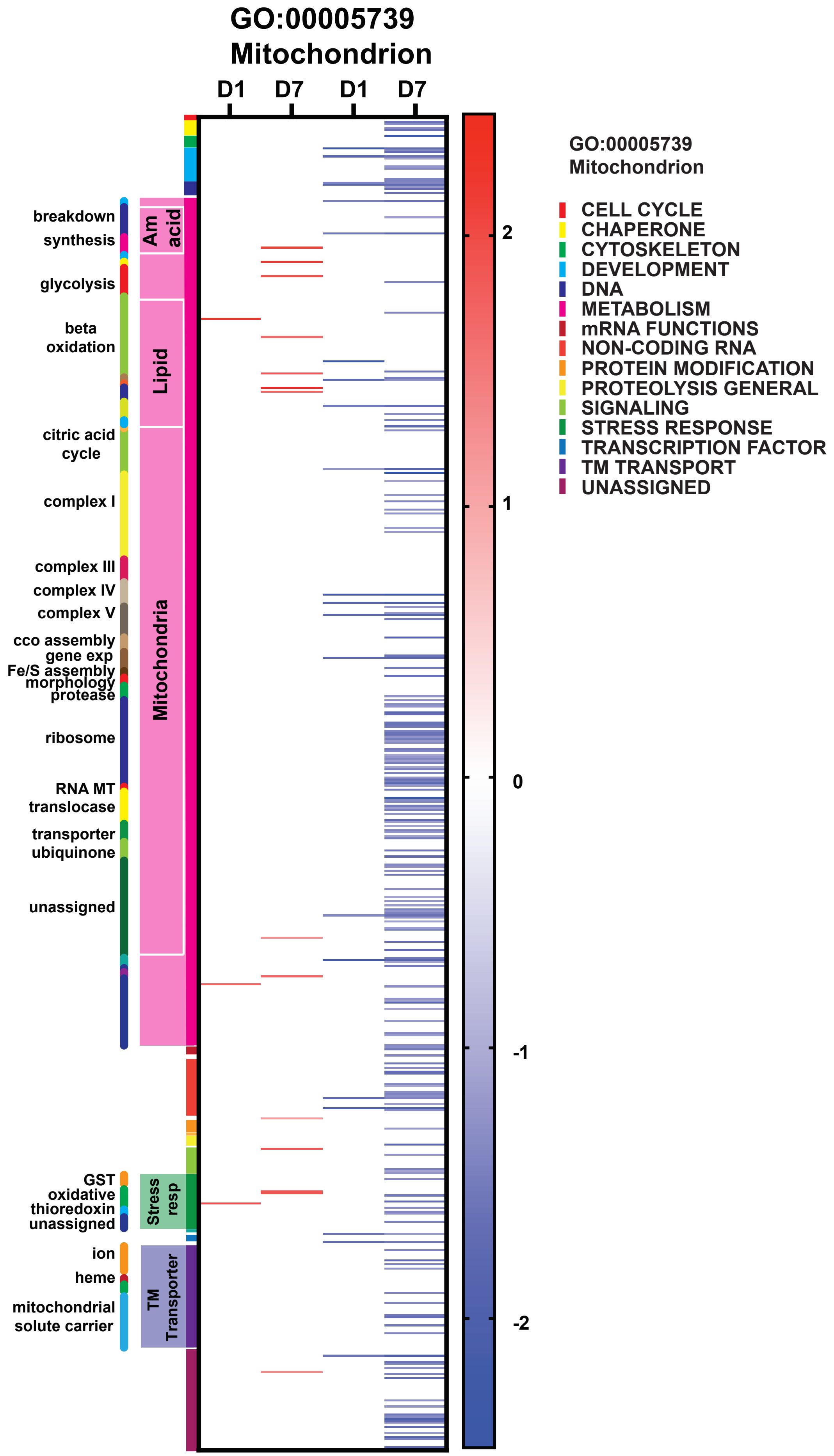

### Supplemental Figure 4

Fig S4
Munden, et al.

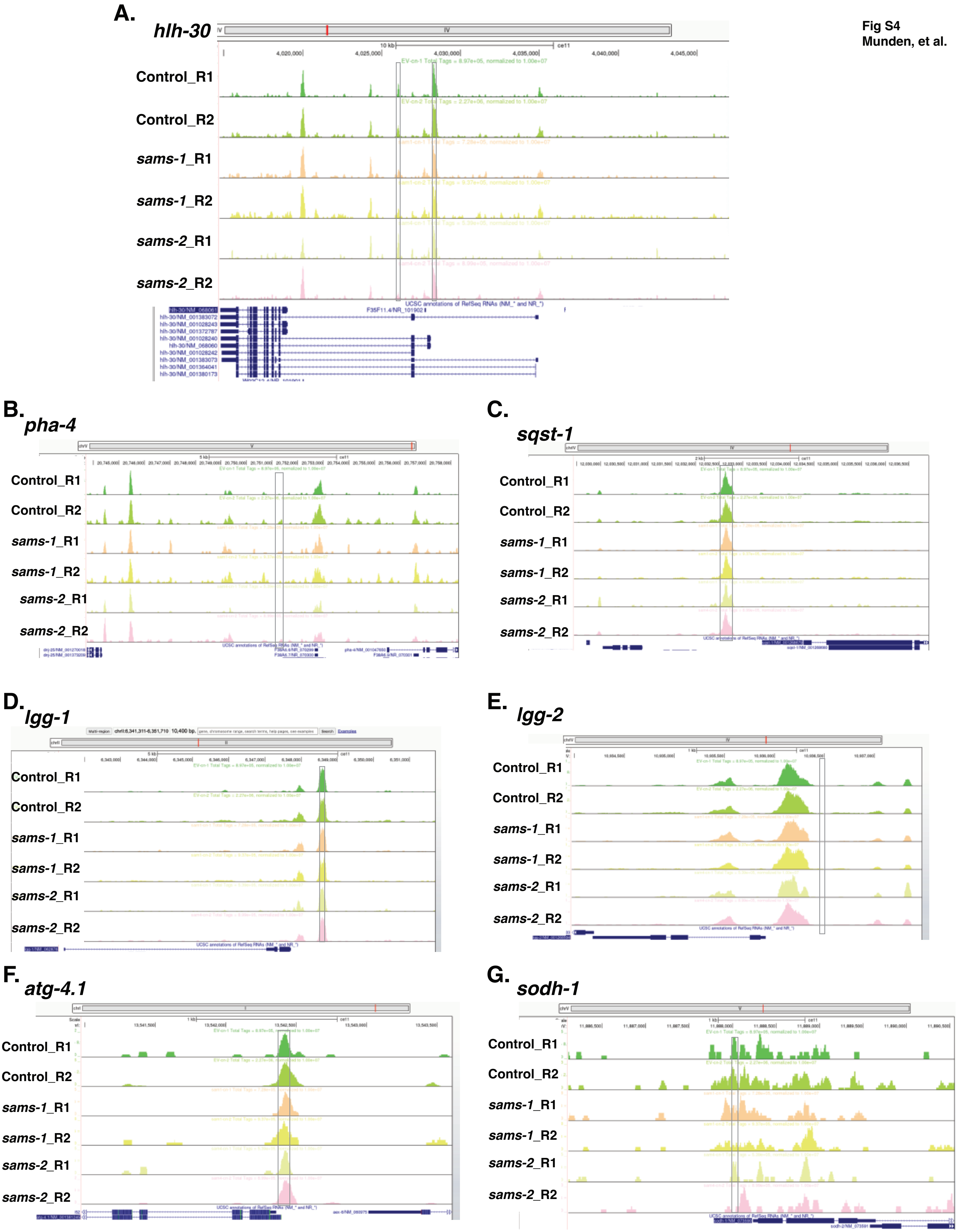

### Supplemental Figure 5

Figure S5
Munden, et al.

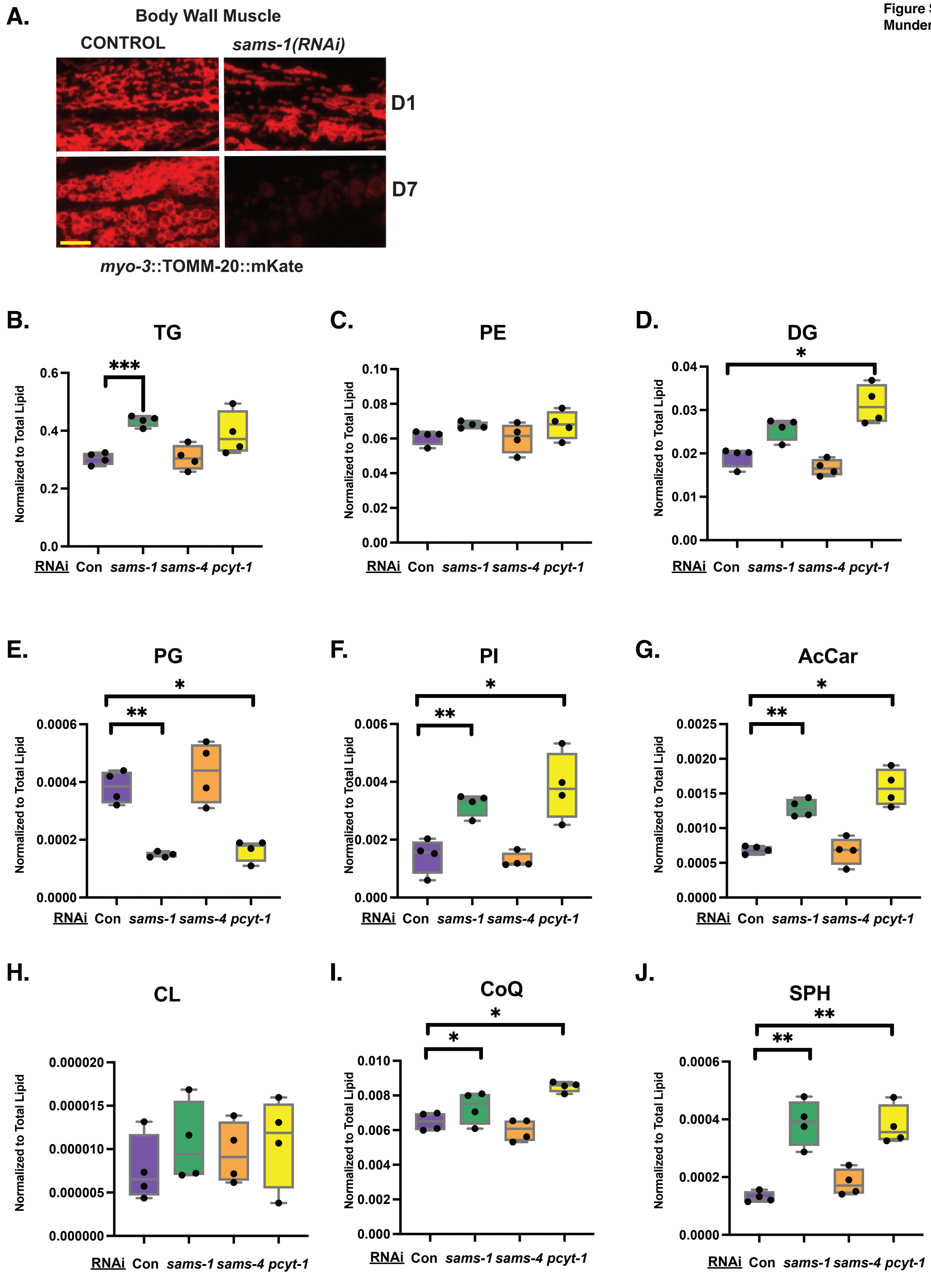

### Supplemental Figure 6

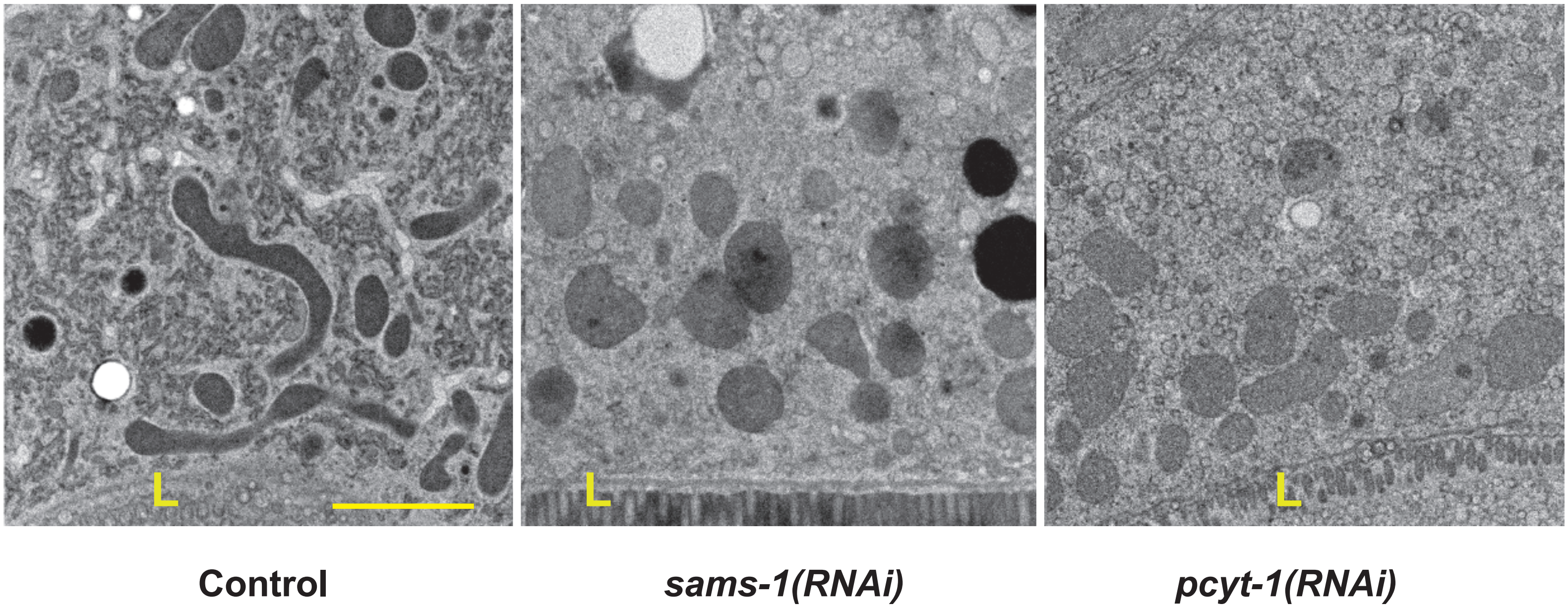

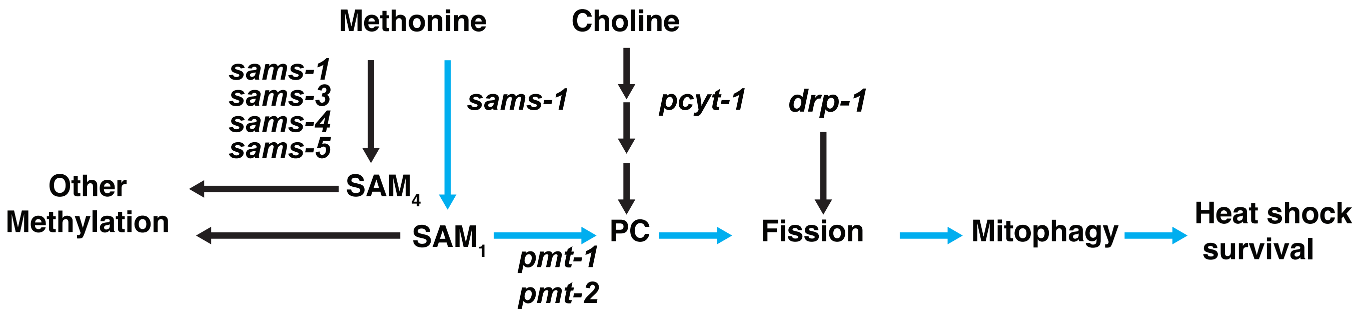
